# Modeling PPRV pathogenesis in mice to assess the contribution of innate cells and anti-viral T cells

**DOI:** 10.1101/2020.10.27.358309

**Authors:** Yashu Sharma, Roman Sarkar, Ayush Jain, Sudhakar Singh, Chander Shekhar, ChandraSekar Shanmugam, Muthuchelvan Dhanavelu, Prabhakar Tembure, Rajeev Kaul, Sharvan Sehrawat

**Affiliations:** Department of Biological Sciences, Indian Institute of Science Education and Research Mohali, Sector 81, SAS Nagar Knowledge City, PO Manauli Mohali 140306 Punjab India; Division of Virology, Indian Veterinary Research Institute, Mukteshwar UK; Department of Veterinary Microbiology, Nagpur Veterinary College, Nagpur MH 440001; Department of Microbiology, University of Delhi South Campus, Benito Zuarez Road New Delhi 110021

## Abstract

We demonstrate a rapid induction of type I IFN response in PPRV stimulated cells and the susceptibility of mice, genetically ablated of interferon response, to PPRV infection. Following PPRV infection, IFNR KO mice gradually reduced their body weights and succumbed to the infection within 10 days. While the infecting inoculum size did not alter the outcome of infection, the nature of the induced disease was qualitatively different. Immunopathological lesions were characterized by the expansion and infiltration of innate immune cells distinctly evident at the lower infecting dose of PPRV infection. The replicating virus particles as well as the viral antigens were abundant in most of the critical organs of PPRV infected IFNR KO mice. Neutrophils and macrophages transported the replicating virus to central nervous system and contributed to pathology while the NK cells and T cells were protective against the virus. Using an array of fluorescently labeled H-2K^b^ tetramers PPRV specific CD8^+^ T cells responses were identified and measured in the infected as well as the peptide immunized mice. Our study therefore established and employed a laboratory animal model for investigating PPRV pathogenesis and the contribution of virus specific CD8^+^ T cells during the virus infection to pave the way for elucidating protective or pathological roles of immune cells during PPRV infection.

**Importance:** We developed a laboratory animal model for investigating the pathogenesis and immunity induced by PPRV. IFNR KO animals succumbed to the infection irrespective of the dose and the route of infection. Neutrophils and macrophages served as the Trojan horse and helped transport the virus to CNS to cause encephalitis while the NK cells and CD8^+^ T cells provided the protection against PPRV infection. We additionally identified class I restricted immunogenic epitopes of PPRV in C57BL/6 mice. Our study therefore paves the way for an optimal utilization of this model to unravel PPRV pathogenesis and assessing the host correlates of protection.

## Introduction

Peste des petits ruminants virus (PPRV) causes high mortality in herds of small ruminants such as sheep and goats and is responsible for major economic losses to livestock sector in developing countries (1–4). PPRV is a negative sense, single stranded enveloped virus of paramyxoviridae family that also include other members that cause debilitating diseases in animals as well as humans. These include Rinderpest virus (RPV) and Canine distemper virus (CDV) of animals and measles virus (MeV) and mumps virus (MuV) of humans (5, 6). PPRV genomes encode for six structural proteins i.e., nucleocapsid (N), phosphoprotein (P), matrix (M), fusion (F), hemagglutinin (H) and polymerase (L) and two nonstructural proteins, C and V (5). The protective and pathological mechanisms activated by the virus as well as the roles of immune cells in its pathogenesis have not yet been adequately elucidated that is primarily attributed to the unavailability of a laboratory animal model. We therefore investigated PPRV pathogenesis in a more accessible vertebrate laboratory animal model to better understand immunity and immunopathology induced by PPRV.

Currently, a live attenuated prophylactic vaccine against PPRV is used in small ruminants. While the vaccine induces a lasting immunity, a transient immunosuppression is usually evident in vaccinated animals that could enhance their susceptibility to heterologous infections (3, 5). Therefore, it is imperative to study the contribution of cellular and molecular mediators induced by the virus in infected animals to better understand its pathogenesis and help devise improved vaccination strategies should a need arise. This is even more relevant for the contemporary animal health care system as the extensive efforts are made to eradicate PPRV globally. That an intensive vaccination program could help eradicate PPRV is bolstered by the success achieved in eradicating a related RPV. Therefore, an accessible laboratory animal model for elucidating PPRV pathogenesis is likely to improve our understanding of the induced molecular and cellular mediators. Similarly such a model would shed light on the host correlates of protection.

We demonstrate a rapid induction of type I IFNs (α and β) as well as IFN-γ response in PPRV-stimulated immune cells but the kinetics of response varied depending on the cell types and the dose of the stimulating virus. Mice genetically deficient for IFN response (AG129) succumbed to the infection within ten days irrespective of the dose of inoculum or the route of PPRV infection. The inoculum size altered the pathology qualitatively. A lower infecting dose of the virus induced predominantly an immunopathological response in mice infected with PPRV via intranasal route. The replicating PPRV as well as its antigens were detected in most of the analyzed organs. Innate immune cells such as neutrophils and macrophages likely transported the replicating virus to CNS and elsewhere. A reconstitution of IFNR KO mice with wild type CD8^+^ T cells conferred a survival advantage during infection, a suggestion for their critical role in the PPRV control. We also identified immunogenic class I (H-2K^b^) restricted epitopes of PPRV in mice using epitope prediction tools and demonstrated their immunogenicity *ex vivo* and *in vivo*. Therefore, our study established a laboratory animal model that could be valuable for understanding the immunological and virological parameters of morbilivirus induced diseases.

## Materials and Methods

### Virus and cells

PPRV vaccine strain Sungri/96 was used for all the *ex vivo* and *in vivo* experiments. The virus was cultured, harvested and titrated using Vero cells and stored at −80°C till further use as described earlier (4, 7). The infecting dose of the virus was calculated as TCID_50_ values by a previously described method (8). RAW264.7 cells were cultured in complete RPMI medium supplemented with 10% FBS and penicillin-streptomycin in a humidified CO_2_ incubator.

### Antibodies and other biological reagents

Antibodies used in this study were purchased from BD Biosciences, Tonbo biosciences, BioLegend, and eBiosciences. The antibodies used were against CD4 (clone GK1.5), purified CD16/32 (Clone 2.4G2), CD11b (clone M1/70), Gr1 (clone RB6-8C5), F4/80 (clone T45-2342), CD8 (clone 53-6.7), H2K^b^ (clone AF6 88.5), mouse IgG, CD45.1 (clone A20), CD45.2 (clone 1O4), CXCR3 (clone 173), CD44 (clone IM7), CD62L (clone MEL 14) and CD45 (Clone 30-F11). All the antibodies were diluted in FACS buffer. Other reagents such as DMEM, RPMI 1640, and penicillin-streptomycin antibiotic were purchased from Lonza. Trypsin, SYBR Green and propidium iodide were obtained from Life Technologies. H&E was from HiPrep, M-CSF was from Peprotech and OCT compound was obtained from Fisher HealthCare. FBS, p-nitrophenol phosphate and Freund’s complete and incomplete adjuvant were procured from Sigma-Aldrich.

### Generation of bone marrow derived macrophages (BMDMs)

Long bones were collected from sacrificed C57BL/6 mice and sterilized in 70% alcohol. Bone marrows were removed to prepare single cell suspension as described earlier (9). RBCs present in the bone marrow cells were lysed and the bone marrow cells were cultured in 48 well plates (1×10^6^ cells/well) in the presence of 10ng/ml M-CSF for 7 days. Media was changed after every two days. The cells were cultured in 10% RPMI (Gibco◻BRL, Rockville, MD, USA) supplemented penicillin (100U/mL) and streptomycin (100μg/mL). After 7 days, cells were collected, washed and stained for F4/80 positive and CD11c negative population for phenotypic characterization and for performing further experiments.

### Measuring type I and II IFN response in PPRV pulsed macrophage cell line and primary BMDM cells

RAW macrophages and BMDMs were pulsed with PPRV at multiplicity of infection (MOI) of 1 and 10 to measure the kinetics of type I IFNs (α and β) and IFN-γ response by qualitative real time polymerase chain reaction (RT-PCR). Murine macrophages and RAW cells pulsed with replicating PPRV or the inactivated virus and samples were collected at 15 min, 30 min, 1hr and 6hr. The cells were processed for isolating mRNA at different time points. The mRNA was converted into cDNA using a first strand synthesis kit (Verso cDNA synthesis kit, Thermo Fisher Scientific). The expression of hypoxanthine phosphoribosyltransferase 1 (HPRT 1) gene served as an internal control. The relative expression for each gene was calculated by using Δ(ΔCt) method.

### Infection of mice with PPRV

All the experiments involving animal experiments were performed strictly in accordance with the protocol approved by the Institutional Animal Ethics Committee, IISER Mohali, constituted under the aegis of committee for the purpose of control and supervision of experiments on animals (CPCSEA). IFNR KO (AG129) and congenic C57BL/6 mice (CD45.1 and CD45.2) were used for *in vivo* experiments. Animals were infected using intraperitoneal (i.p) or intranasal (i.n) routes with the indicated doses of PPRV. Different physiological parameters such as body weight, body temperature, behavior and the mortality pattern were measured in different groups of animals. For most of the experiments, the animals were sacrificed at the termination of experiments when the body weight for any of the groups dropped by more than 20%. Different lymphoid and non-lymphoid organs were collected to detect the replicating virus, viral antigens, and performing the cellular analysis in lymphoid organs such as spleen, mediastinal LNs as well as non-lymphoid organs such as bronchoalveolar lavage (BAL), lungs and brain tissues. Before collecting organs from different groups of mice, a heart perfusion with 20 ml of PBS was performed to remove any contaminating cells of blood from the collected organs.

### Reconstitution of IFNR KO mice with T cells to measure their anti-PPRV functions

In order to measure the protective ability of immune cells, the graded doses of MACS purified CD4^+^ and CD8^+^ T cells from C57BL/6 WT mice were adoptively transferred in sex matched IFNR KO animals. The recipient animals were subsequently infected with PPRV. Mice not transferred with any cells served as the control. Recipient animals were observed for their body wight and survival. At the termination of experiments, lymphoid organs of animals were collected for cellular analysis.

### Cell purification and adoptive transfer

The different subsets of T cells and innate immune cells such as macrophages, neutrophils, dendritic cells were isolated from C57BL/6 mice either by magnetic cell sorting kits or by FACS sorting. The sorted cells were collected at low temperature in the complete RPMI. The cells were pulsed with PPRV for one hour. After extensive washings, the cells were counted and the indicated numbers of cells were transferred i.v in IFNR KO mice.

### Bioinformatic analysis to predict immunogenic peptides of PPRV

Amino acid sequences of PPRV structural proteins were retrieved from National Centre for Biotechnology Information (NCBI) database in FASTA Formats. The immunogenic peptides for one of the class I MHC molecules of C57BL/6 mice (H-2K^b^), were predicted using immune epitope data and analysis resource (IEDB). The software uses artificial network (ANN) and stabilized matrix method (SMM). The percentile rank of <2 and IC_50_ values were selected for the prediction. A low percentile rank and the lower IC_50_ values of <50nM indicated high affinity binders. The peptides with IC_50_ values between 50 and <500nM were considered as intermediate affinity binders while with values >500nM were considered as the low affinity binders (10). Additional parameter for selecting peptides was their immunogenicity scores (11,12). Additionally a percentile rank for the predicted peptides was generated by comparing IC_50_ values of predicted epitopes against a set of random peptides using SWISSPROT database. The selected peptides were commercially synthesized and obtained from GL Biochem. The purity of the synthetic peptides was greater than 90%.

### Molecular Docking analysis

For predicting the binding affinities of different peptides for class I MHC molecules (H-2K^b^ and Caprine Leucocyte antigen, CLA1), molecular docking analyses were performed using HPEPDOCK-web server. Default parameters were used for all the docking experiments as described elsewhere (13). HPEPDOCK server uses a hierarchical algorithm, MODPEP program for a blind protein-peptide docking and generates an ensemble of peptide conformation by considering flexibility conformation in the respective peptide and the PDB File 1S7Q for the homology modeling with H-2K^b^ protein. To test the efficiency of docking algorithm, a known immunogenic 9-mer peptide derived from Sendai E virus (SEV) nucleoprotein (FAPGNYPAL) was used for docking with H-2K^b^. Since the outcome of docking could be dictated by a potentially problematic algorithm that overemphasize the numbers of interactions rather than the conformation, we referred to the solved crystal structure of a nonameric peptide (SEV-9) with H-2K^b^ to better predict the results from docking analyses. We then compared the energy parameters of the best-selected structures among different PPRV peptides docked with H-2K^b^. To further refine and define the interacting residues both quantitatively and qualitatively, we used molecular modeling program UCSF Chimera for binding analysis (14). As the goal of such experiments was to explore the translational value of such peptides in small ruminants, we superimposed H-2K^b^ with goat class I MHC (CLA-1) molecule at 1.5A RMSD (root mean square distance) using the tool, Matchmaker, available in the UCSF Chimera. Similarly docking studies were done for CLA-I with different PPRV peptides and the representative docked structures were generated using UCSF Chimera. A comparative analysis between docking scores of H-2K^b^ and CLA-I for the similar peptides was also performed.

### Class I MHC stabilization assays

The stabilization of class I MHC by each peptide was measured using both cellular and acellular assays. TAP deficient murine T cell lymphoma cells (RMA/s cells) were used for determining the peptide induced surface stabilization of class I MHC molecule by flow cytometric analysis as described earlier (15). RMA/s cells were maintained in RPMI (Gibco◻BRL, Rockville, MD, USA) supplemented with 10% fetal bovine serum (FBS), penicillin (100U/mL) and streptomycin (100μg/mL). 2×10^5^ cells were serum starved for 3 hrs at 37°C and subsequently pulsed with the respective peptides to induce their surface class I MHC (H-2K^b^) stabilization. Graded doses of peptides were added in serum free RPMI followed in which cells cells were incubated at 37°C for 7 hrs. The cells were then washed with PBS and stained with anti anti-H-2K^b^-FITC antibody. Live and dead cells were differentiated using propidium iodide (PI) staining. The cells were analysed by FACS Accuri flow cytometer (BD Biosciences, Breda, The Netherlands). The data is represented as percent positive cells or the mean fluorescence intensities (MFI) for the expression of H-2K^b^. EC_50_ value for high affinity peptides were then calculated.

An acellular assay was also used for determining the class I MHC stabilization as described elsewhere (16). Briefly, ELISA plates were coated with streptavidin overnight at 4°C. Subsequently, H-2K^b^ monomers were added to the plates. The monomers were generated by refolding a UV photocleavable ligand, (FAPG(Anp)YPAL), β2 microglobulin and H-2K^b^ heavy chain followed by their biotinylation as described earlier (15, 17). The unbound H-2K^b^ monomers were washed and the control and PPRV peptides were added to the identified wells in the plates. The plates were then exposed to UV radiations at 365nM for 30 min to achieve the displacement of UV ligand with respective peptides. The efficiency of exchange was measured by probing the washed plates by adding anti-β2 microglobulin antibody. Subsequently, a mouse anti-IgG antibody enzyme conjugate (1:10000) was added after washing the plates. Thereafter, substrate, (p-nitrophenol phosphate (1mg/ml) was added for its conversion into a chromogenic product. The stop solution was used to block the reaction and the plates were measured for absorbance at 405 nm. Positive and negative controls were also included in the study (18).

### PPRV infection and peptide immunization of mice for PPRV specific CD8^+^ T cell analysis

In order to determine the immunogenicity of predicted peptides in vivo, we performed two types of experiments. Throughout the manuscript, plaque forming units (PFU) and focal forming units (FFU) are used interchangeably as clear plaques are not observed when PPRV in grown in Vero cells. In first set of experiments, WT C57BL/6 mice were i.p. infected with a high dose of PPRV (5×10^6^ PFU). After seven days a second dose of PPRV was given to animals to boost responses. The analysis of the expanded cells was performed three days later by measuring the frequencies PPRV-peptide specific CD8^+^ T cells by tetramer staining. In second set of experiments, C57BL/6 mice were immunized either with the cocktail of peptide (AILTFLFLL, FMYLFLLGV, FSAGAYPLL and IGLVRDFGL) each with 5μg/mouse in complete Freund’s adjuvant subcutaneously. After two weeks, a second injection of the same concentration was administered but emulsified in the incomplete Freund’s adjuvant. Three days later the frequencies of peptide specific CD8^+^ T cells were analyzed by tetramer staining of PBMCs.

### Isolation of inflammatory cells from brain tissues and lungs

Inflammatory cells were isolated from the brains and lungs of IFNR KO mice by using protocol as described earlier (19). Briefly, brain tissues were minced into 3-4 mm pieces with a sterile scalpel or scissors under complete aseptic conditions. Washing was done 4-5 times with 10mM PBS. Extra supernatant was removed from tissue pieces container kept on ice. 0.25% trypsin was added to samples followed by their incubation at 4°C for 16 hrs. Then, trypsin was discarded and incubation was done at 37°C for 30 min. Complete RPMI was added to prepare single cell suspension. After washing with PBS, cells were used further for experiments.

### Flow cytometry for cellular analysis

Different lymphoid and non-lymphoid organs were collected from infected or immunized mice. The single cell suspensions were prepared from collected organs for cellular analysis. The cells were stained using indicated fluorescent labeled antibodies at cold temperature for 30 minutes. Fc block was done before surface staining. Stained cells were acquired by FACS Accuri or BD FACS Aria fusion. The analysis of the data was performed by Flow Jo software.

### Fluorescent microscopy and histological analysis

The organs collected from infected and control animals were stored in OCT compound. Tissue sections of 6◻m were cut and dried. The dried sections were first blocked with anti-CD16/CD32 antibodies followed by incubation with anti-PPRV H and N monoclonal antibodies. Anti mouse FITC IgG antibodies were then used as secondary antibodies. The sections were analyzed using a fluorescent microscope and the images were generated by Image J software. Similarly, the tissue section from brain tissues were dried and stained with Hematoxylin and Eosin Y. The sections were analyzed by Leica DMi8 microscope as described earlier (20).

### Statistical analysis

Statistics was applied to the data and analysed by using ANOVA, Student t test or the Gehan-Breslow-Wilcoxon test as indicated in the respective figures. Graph Pad Prism v5.03 was used for such analysis. The level of significance was determined as P < 0.05 *, P < 0.01 **, P < 0.001***, P < 0.0001****.

## Results

### PPRV infection induces a rapid interferon response

Interferon response constitutes the first line of an anti-viral defense mechanism. We therefore measured the expression of both type I (α, β) and type II (γ) IFN response in a murine macrophage cell line (RAW macrophages) and the primary BMDMs that were stimulated with a low (1MOI) and high (10MOI) dose of PPRV. A low dose of PPRV as compared to the high dose induced significantly more IFNαin RAW macrophages at 1hr post stimulation (Fig 1A). However, a reverse trend as well as an early induction was observed in BMDMs (Fig 1D). Furthermore, the overall expression levels of IFNα were approximately 100 fold more in the primary BMDMs as compared to those in RAW macrophages (Fig 1A and D). Interestingly, we observed a very rapid induction of IFNα within 15 min of stimulation by primary BMDMs but its expression was evident in RAW macrophages only after 1 hour (Fig 1A and D). Similarly IFNβ expression was only detectable at significant levels in PPRV stimulated RAW macrophages after 1hour (Fig 1B). In stimulated BMDMs, the message of IFNβ was evident albeit at lower levels as compared to that of IFNα (Fig1E). IFNγ was induced at a low infecting dose of PPRV in RAW cells but the primary BMDMs expressed it in significant levels only at the high MOI of PPRV (Fig 1C and F). The heat inactivated PPRV induced the production of both type I and type II IFNs but to a much lesser extent and that too in the early stages of stimulation (Fig S1). These results might suggest that not only the viral genome or its replication intermediates but also some of the PPRV proteins could serve as the PAMPs for inducing IFN responses.

**Figure 1.**
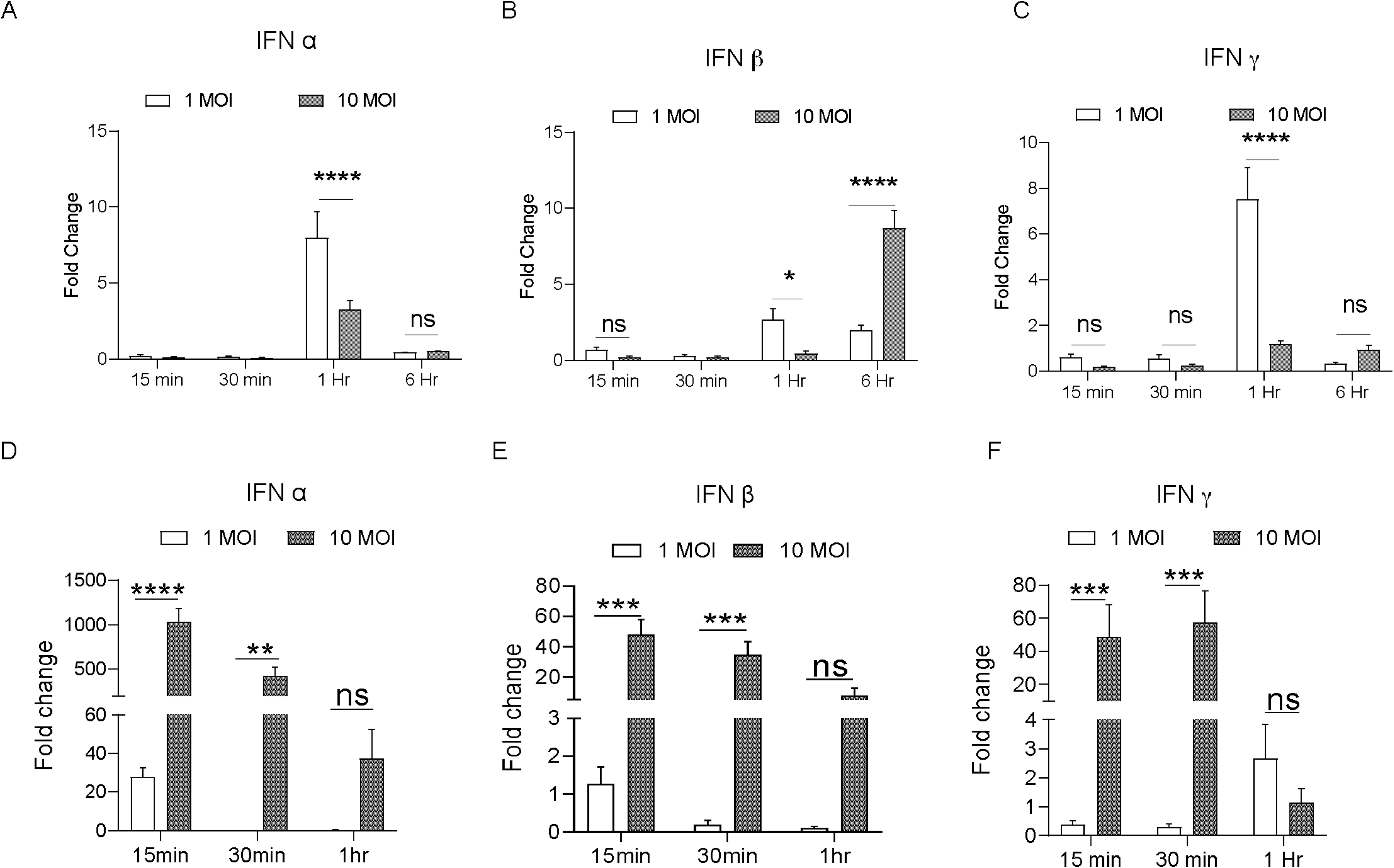
PPRV pulsed murine macrophages mount a rapid IFN response. Murine macrophages (RAW macrophages and primary bone marrow derived macrophages) were pulsed with PPRV at low (1:1) and high (1:10) multiplicity of infections (MOI) to measure IFNα, IFNβ and IFNγ response. The PPRV exposed cells were collected at different times to isolate total RNA. cDNA synthesized was measured for the expression of different IFNs by RT-qPCR. A-C. Fold change in the expression of IFNα, β and γ is shown by bar diagrams at 15min, 30min, 1hr and 6hr. D-F. BMDMs similarly analyzed for the expression of IFNα, β and γ at indicated time points. Bar diagrams show fold change in the expression of IFNs. The experiments were repeated six times. One way ANOVA test was used for determining the statistical significance and p values are represented as following; p < 0.05 *, p < 0.01 **, p < 0.001***.

Our results demonstrated that both α and β IFNs were induced in PPRV stimulated cells but the overall expression was dependent on the infecting dose as well as the nature of responding cells. The expression profile of type I interferon (IFNα and IFNβ) also indicated a dichotomy in their function with the IFNα being induced rapidly and in high concentrations but IFNβ was produced later on. Such a response pattern could diversify the IFN response in providing an antiviral state.

### Mice genetically depleted of IFNRs are susceptible to PPRV infection

Having demonstrated a rapid induction of IFN response in the PPRV pulsed macrophages; we tested whether or not the mice, unable to mount IFN response because of genetic ablation of the signaling receptors, are susceptible to PPRV. Different doses of PPRV (1, 10^2,^ 10^3^ and 10^4^ PFU) were i.p inoculated into IFNR KO mice and the disease progression was monitored (Fig 2A). We first measured the survival of PPRV infected animals up to eight dpi. All the infected animals succumbed to the infection albeit survival duration was dependent on the initial inoculum (Fig 2B). Accordingly, animals infected with the high dose died earlier as compared to those infected with the lower dose of PPRV (Fig 2B). We then measured other physiological parameters and body weights in the infected animals (Fig 2C). All the infected animals gradually lost their body weights, developed encephalitic lesions and became hypothermic (Fig 2C and data not shown). By 7dpi, all the animals succumbed to the infection irrespective of the doses of virus inoculum used (Fig 2C). In similar experiments, PPRV infected WT mice remained refractory to the PPRV induced disease as no clinical signs were observed even in those infected with the high infecting dose (5×10^6^ PFU) (Fig 2B and C, S6B). The results, therefore, underscored the critical role of IFN signaling in providing an early defense against PPRV infection in mice.

**Figure 2.**
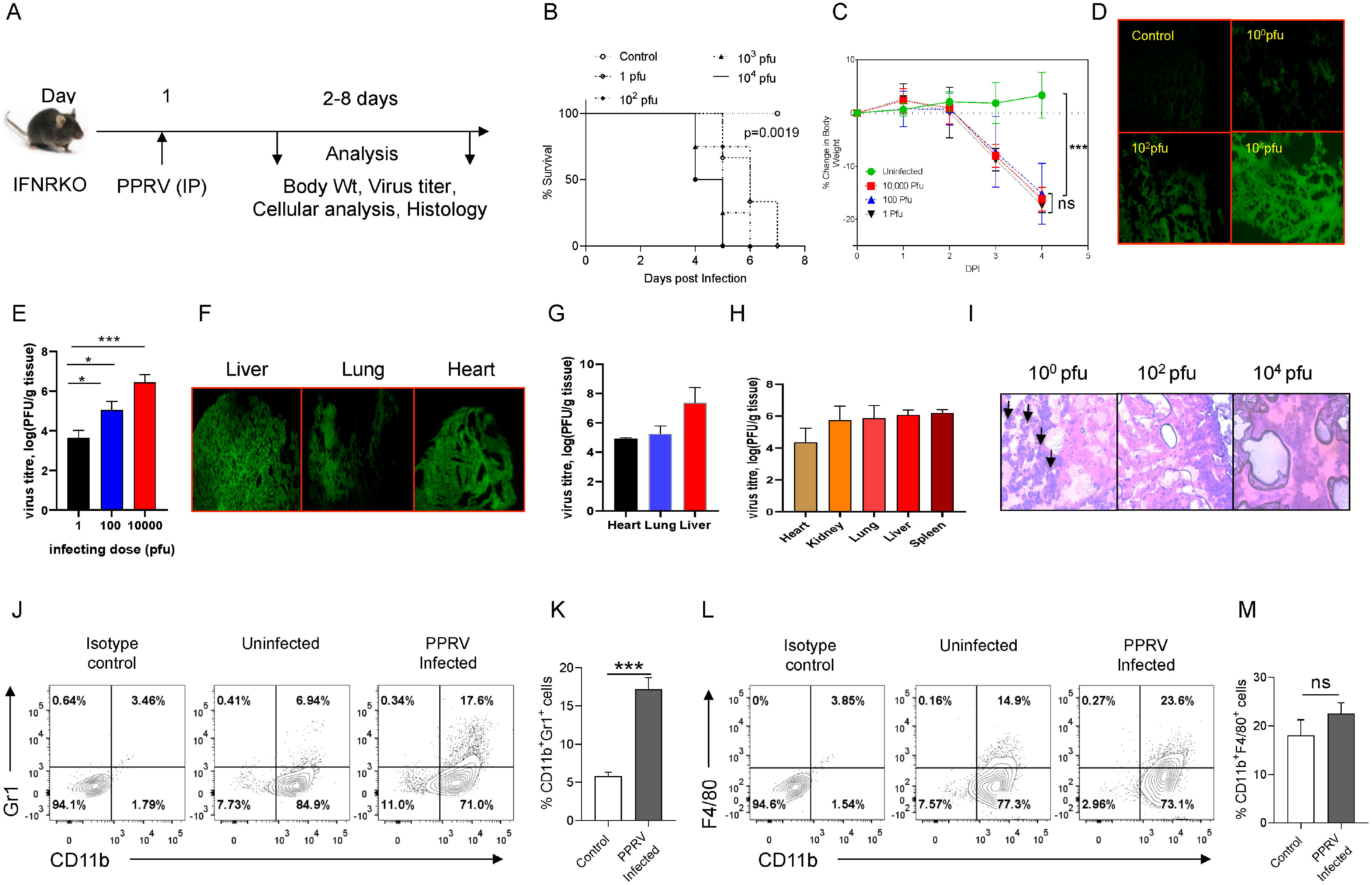
IFNR KO mice are susceptible to PPRV infection. A. The schematic of experiments is shown. IFNR KO mice were i.p. infected with indicated doses of PPRV and the survival analysis (B) and percentage change in body weight of mice (C) from each group is shown. The level of statistical significance was determined by Gehan-Breslow-Wilcoxon test and one-way ANOVA, respectively. Each data point in the graph represents the average percent change in body weight at the indicated dpi. Control animals were not infected with PPRV. D. Fluorescent microscopic images show the presence of PPRV antigens in brain tissues of the infected IFNR KO mice as detected by anti-PPRV (H) mAbs. E. Bar diagrams show PPRV titres as represented by log10 PFU/g of tissue when measured by plaque forming assays using brain tissues of i.p., infected IFNR KO mice administered with varying doses. F-G. IFNR KO mice were i.p. infected with 1.5×10^6^ PFU of PPRV and different organs were collected 4dpi to measure the presence of viral antigens (F) and replicating virus (G) in different organs. F. Fluorescent microscopic images show the presence of virus antigens in brain tissues of PPRV infected IFNR KO mice as detected by anti-PPRV (H) protein mAbs. G. Virus titers as represented by log10 PFU/g of tissue in heart, lung and liver is shown. Data represents the mean ± SD of three replicates of tissue samples. H. Virus titers as represented by log10 PFU/g of tissue are shown in different organs of IFNR KO mice infected with 1PFU of PPRV. I. Histopathological changes in virus infected brain sections are shown. The experiments were repeated three times with similar results. B-I. One way ANOVA test was used for determining the statistical significance values and are represented as following; p < 0.05 *, p < 0.01 **, p < 0.001***. J-M. Innate immune cells were analyzed in the brain tissues of PPRV infected IFNR KO mice. The single cell suspensions prepared from brain samples were stained with neutrophils (CD11b^+^Gr1^+^) and macrophages (CD11b^+^F4/80^+^). J. Representative FACS plots show the frequencies of CD11b^+^Gr1^+^ cells in control and PPRV infected mice. K. Bar diagrams show the cumulative data. L. Representative FACS plots show the frequencies of CD11b^+^F4/80^+^ cells in control and PPRV infected mice. M. Bar diagrams show the cumulative data. The experiments were repeated two times. Student t test were used form determining the significance levels in control and infected groups. p < 0.05 *, p < 0.01 **, p < 0.001 ***.

PPRV infected mice expressed encephalitic lesions. We, therefore, measured the presence of PPRV antigens in the brain tissue sections by fluorescent microscopy using anti-PPRV (H) monoclonal antibodies. We also measured the replicating virus particles in brain tissue homogenates by performing plaque-forming assays. Fluorescent microscopic images of brain tissue sections from the infected animals revealed an abundance of viral antigens particularly when the animals were infected with the high doses of the virus inoculum (Fig 2D). A dose dependent increase in the PPRV loads was recorded in the homogenized brain tissues (Fig 2E). Accordingly, the replicating virus titers were 3.6 ± 0.3, 5.0 ± 0.4 and 6.4 ± 0.3 log_10_/g of brain tissues at the infecting dose of 1, 10^2^ and 10^4^ PFU, respectively (Fig 2E). We also collected different organs such as lungs, livers, hearts, brains and kidneys of mice infected with 1 or 1.5×10^6^ PFU of PPRV to measure the virus loads as well as to detect the presence of viral antigens (Fig 2F-H). The fluorescent microscopic images showed the presence of PPRV antigens in the tissue sections of liver, lung and heart of infected animals given 1.5×10^6^ PFU (Fig 2F). The virus titers in liver, lung and heart were 7.3 ± 1.0, 5.2 ± 0.5 and 4.9 ± 0.08 log_10_ PFU/g of tissue, respectively (Fig 2G). The virus load in the animals infected with a low dose of 1 PFU of PPRV were 4.3 ± 0.8, 5.8 ± 0.8, 6.0 ± 0.3, 6.1 ± 0.2 and 5.7 ± 0.8 PFU log_10_/g of tissue in heart, lungs, liver, spleen and kidneys, respectively (Fig 2H). Our results therefore suggested an active replication of PPRV in different organs of mice deficient in IFNs signaling.

IFNR KO mice were susceptible to PPRV infection even at a very low inoculum size (1PFU); we therefore investigated whether or not the infecting dose qualitatively influenced the disease. The brain tissue sections from animals infected with varying doses of PPRV were stained with H&E and the single cell suspensions from brain tissues were analyzed by flow cytometry (Fig 2I). A high dose of PPRV induced swelling of meninges in the infected brain tissues (Fig 2I, right panel). Interestingly, the stained sections of brain tissues from animals infected with low dose of PPRV (1PFU) showed greater leukocytic infiltration as compared to those infected with the high dose (Fig 2I, left panel). We also analyzed the phenotype of leukocytes from the single cell suspension of brain tissues by flow cytometry. The leukocytes (CD45^+^ cells) were analyzed for neutrophils (CD11b^+^Gr1^+^) and macrophages (CD11b^+^F4/80^+^). Significantly higher frequencies of neutrophils were present in the brain samples of PPRV infected mice as compared to the uninfected controls (Fig 2J and K). The frequencies of macrophages also increased but the results were not statistically different in control and infected mice (Fig 2L and M). The increase in the frequencies of innate immune cells further supported the results that the infected mice developed encephalitic lesions.

Many members of genus morbillivirus are neurovirulent and the PPRV induced neurovirulence in naturally infected goat neonates was shown recently (21–23). Therefore, the infectivity of IFNR KO mice by PPRV, their observed neurovirulence as well as immune cells infiltration in infected tissues could suggest that these mice could represent a better accessible model for deciphering molecular and cellular mechanisms occurring during PPRV pathogenesis.

### Innate immune cells are permissive to PPRV infectivity

We established the infectivity of IFNR KO mice and showed the responsiveness of adaptive and innate immune cells. We then measured the immune response in spleens of PPRV infected IFNR KO mice (Fig 3). The frequencies of innate immune cells such as neutrophils (CD11b^+^Gr1^+^) increased by upto 10 fold in PPRV infected mice as compared to controls (Fig 3A and B). Other innate immune cells such as macrophages (CD11b^+^F4/80^+^) and DCs (CD11b^+^CD11c^+^) also showed a significant increase in PPRV infected mice in comparison to control but to a lesser extent as compared to those of neutrophils (Fig 3F, G, K and L). A rapid induction of innate immune cells after PPRV infection and the expression of encephalitic lesions in infected IFNR KO mice even at a low dose of the virus led us to explore the possibility of virus transport to CNS by such cells. We, therefore, measured the infectivity of innate immune cells by PPRV using intracellular staining for the viral proteins. Neutrophils (Gr1^+^), macrophages (F4/80^+^), and DCs (CD11c^+^) from control and the infected animals were measured for the presence of PPRV heamaglutinin (H) and nucleocapsid (N) proteins using monoclonal antibodies (24). In morbilliviruses, N proteins are highly conserved and represent a major component of ribonucleoprotein complex while the H proteins help virus attach to the target cells. A significantly higher frequency of neutrophils isolated from PPRV infected mice showed the presence of H and N viral proteins (Fig 3C-E). Although PPRV proteins were also present in the macrophages and DCs of infected animals but the results were not statistically significant (Fig 3H-J and M-O). Therefore, neutrophils and perhaps other innate immune cells by virtue of their PPRV infectivity might contribute to the virus transport to different tissues.

**Figure 3.**
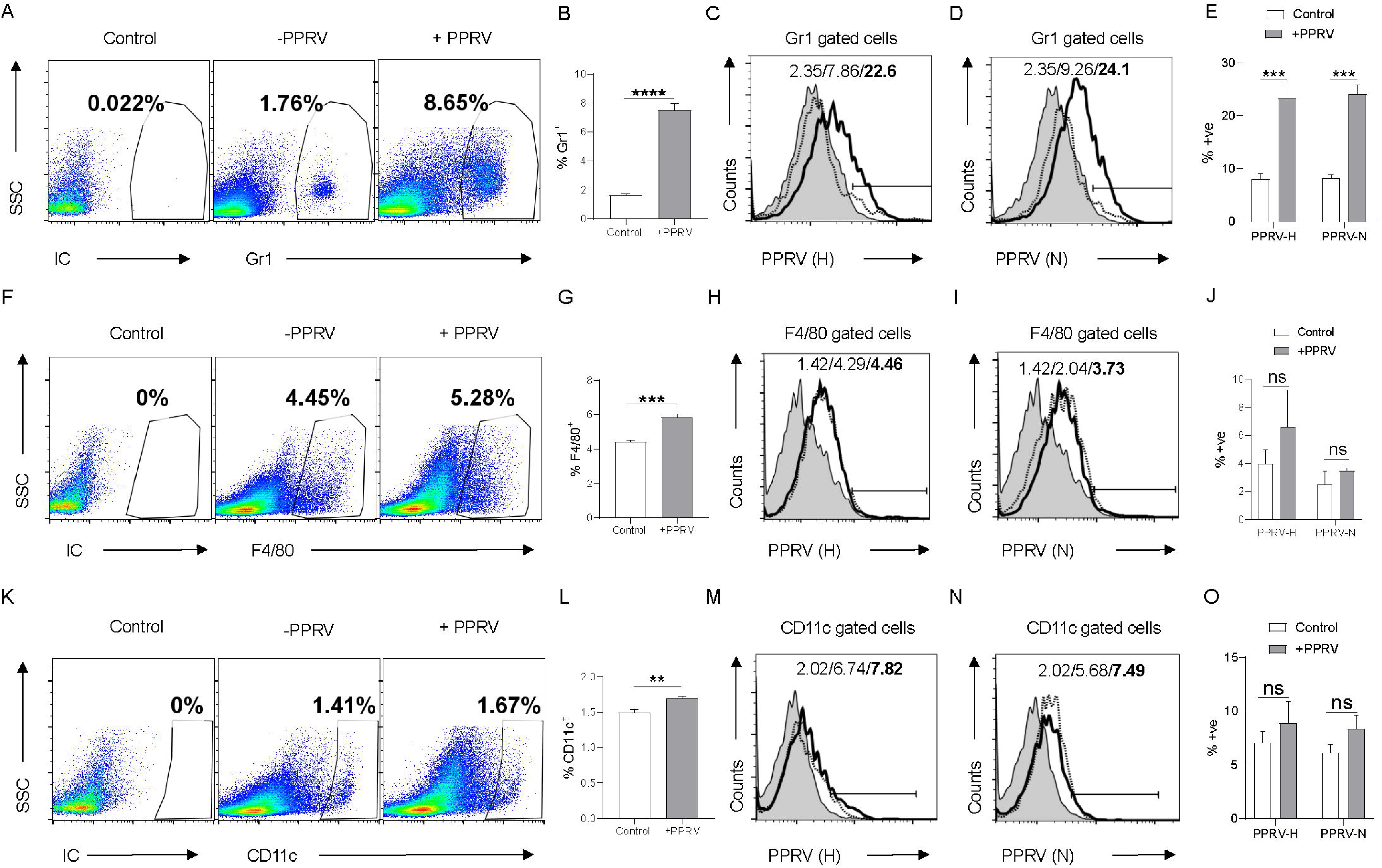
Determining PPRV infectivity of innate immune cells in IFNR KO mice. IFNR KO mice were i.p infected with 100PFU of PPRV and the expansion as well infectivity of innate immune cells was measured in spleen. A. Representative FACS plots show the frequencies of Gr1^+^ cells among live cell gate in control and PPRV infected mice. B. Bar diagrams show cumulative data for the frequencies of Gr1^+^ cells in control and PPRV infected mice. C-E. Intracellular staining was performed as described in the material and methods sections to measure the presence of PPRV antigens in Gr1^+^ cells using anti-PPRV (H) and anti-PPRV (N) protein antibodies. C. Representative overlaid histograms show the frequency of PPRV (H)^+^Gr1^+^ cells. D. Representative histograms show the percentage of PPRV (N)^+^Gr1^+^ cells. E. Bar diagrams show cumulative data as percent positive Gr1^+^PPRV (H)^+^ and PPRV (N)^+^ cells. F. Representative FACS plots show the frequencies of F4/80^+^ cells among live cell gate in control and PPRV infected mice. G. Bar diagrams show cumulative data for the frequencies of F4/80^+^ cells in control and PPRV infected mice. H-J. Intracellular staining was performed as described in the materials and method section to measure the presence of PPRV antigens in Gr1^+^ cells using anti-PPRV H protein and anti-PPRV-N protein mAbs. H. Representative overlaid histograms show the frequency of PPRV (H)^+^F4/80^+^ cells. I. Representative histograms showing the percentage of PPRV (N)^+^F4/80^+^ cells. J. Bar diagrams show the percentage of F4/80^+^ cells expressing PPRV (H) and PPRV (N) proteins. K. Representative FACS plots show the frequencies of CD11c^+^ cells among live cell gate in control and PPRV infected mice. L. Bar diagrams show cumulative data for the frequencies of CD11c^+^ cells in control and PPRV infected mice. M-O. Intracellular staining was performed as described in the material and methods sections to measure the presence of PPRV antigens in CD11c^+^ cells using anti-PPRV H protein and anti-PPRV-N protein antibodies. M. Representative overlaid histograms show the frequency of PPRV (H)^+^CD11c^+^ cells. N. Representative histograms show the percentage of PPRV (N)^+^CD11c^+^ cells. O. Bar diagrams show the percentage of F4/80^+^ cells expressing PPRV (H) and PPRV (N) proteins. The experiments were repeated two times with four animals in each group. Different groups were analyzed by two-way ANOVA using Sidak’s multiple comparison test. Statistical significance values are represented as following; p < 0.05 *, p < 0.01 **, p < 0.001***.

### Innate immune cells transport PPRV to central nervous system

We investigated whether or not innate immune cells help transport PPRV to CNS. PPRV infected innate immune cells were adoptively transferred in congenic mice followed by monitoring the disease progression in recipients (Fig 4A). Neutrophils, macrophages and dendritic cells were FACS sorted from WT congenic mice (CD45.1^+^). The sorted cells were infected with PPRV and after extensive washings; these cells were transferred into sex matched IFNR KO mice (CD45.2^+^) (Fig 4A). We recovered enhanced frequencies of neutrophils (CD45.1^+^CD11b^+^Gr1^+^, ~ 4%) and macrophages (CD45.1^+^CD11b^+^F4/80^+^, ~ 2%) from brain tissues of recipients (Fig 4B and C). Moreover, the transferred PPRV-pulsed cells resulted in a patent infection in recipients with the progression of disease being similar to that observed in PPRV only injected animals (Fig 4D). This data suggested that infected innate immune cells particularly the neutrophils and macrophages could support virus replication and transport PPRV to different organs. We did not recover significantly higher frequencies of donor DCs (CD45.1^+^CD11b^+^CD11c^+^) in the brain tissues of recipient mice but the disease severity was comparable in all the recipients (Fig 4B-D). Several factors could explain these results such as the inability of transferred DCs to directly home to brain tissues or their inefficient proliferation. Nonetheless, such cells might have transferred the virus to other inflammatory cells, which then could have transported it to brain tissues and elsewhere. The observed increased leukocytic infiltration in the brain tissues supported this notion (data not shown). Further analysis of different innate immune cells pulsed with PPRV demonstrated their infectivity as PPRV antigens could be detected in such cells by flow cytometry (Fig 4E and F). These results indicated that the virus could either be internalized by innate immune cells or be associated to surfaces and such cells could transport PPRV to distant sites such as the CNS.

**Figure 4.**
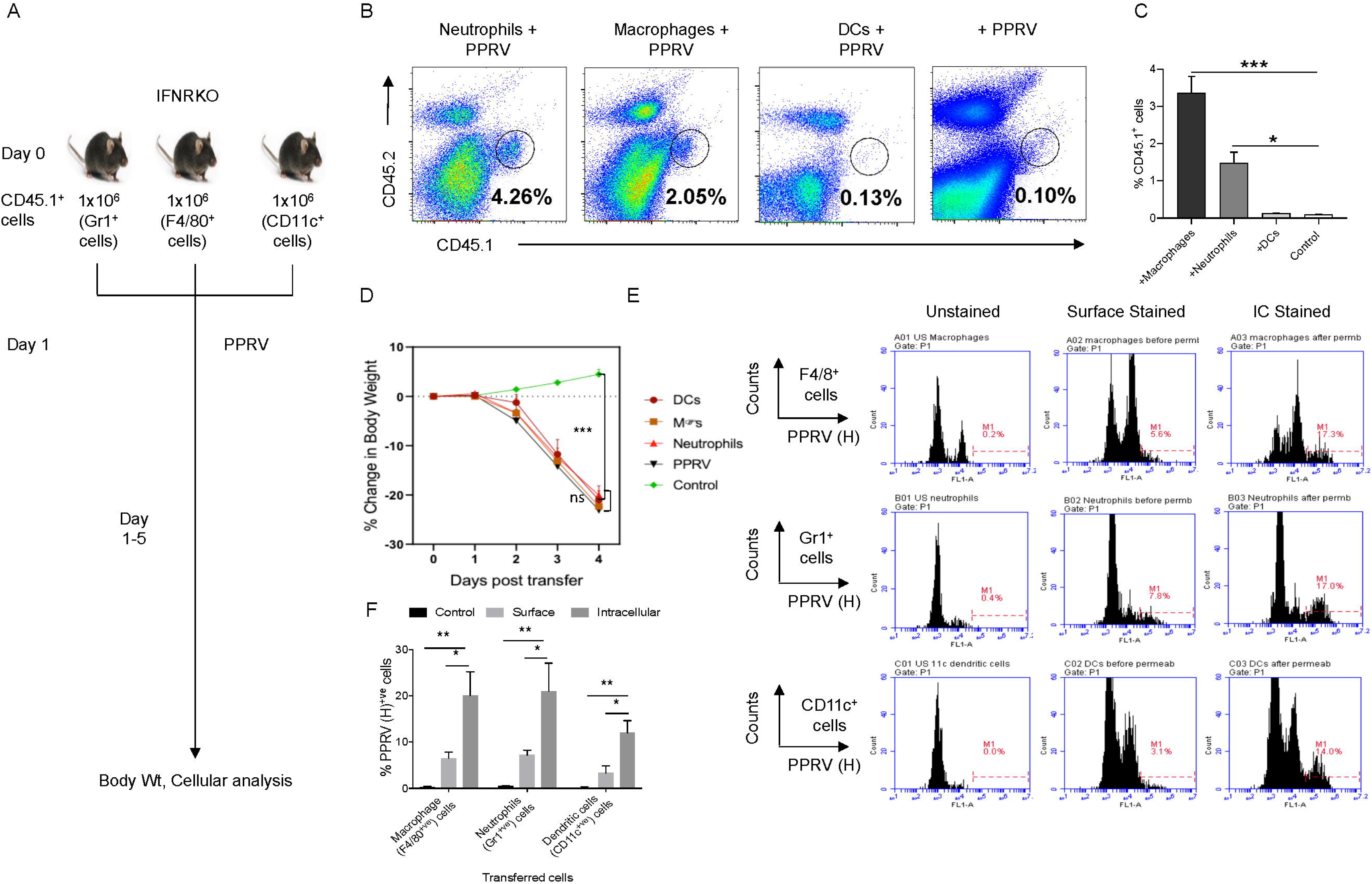
Innate immune cells serve as the carrier of PPRV to brain tissues. FACS sorted neutrophils, macrophages and dendritic cells from CD45.1^+^ C57BL/6 mice were pulsed with PPRV and after washing transferred into CD45.2^+^ IFNR KO mice. At 4dpi, brain tissue staining was done with CD45.1 and CD45.2 markers. A. Representative FACS plots depict the frequencies of donor cells in processed brain tissue suspensions. B. Bar diagrams shown the cumulative frequencies of frequencies expanded donor cells recovered from brain suspensions. For each group of recipients three animals were used and the experiments were performed two times. C. FACS sorted macrophages (F4/80^+ve^), neutrophils (Gr1^+ve^) and dendritic cells (CD11c^+ve^) from CD45.1^+^ C57BL/6 mice were pulsed with PPRV for 1hr and transferred in IFNR KO mice. The recipients were monitored for their body weights and morbidity. D-E. FACS sorted macrophages (F4/80^+ve^), neutrophils (Gr1^+ve^) and dendritic cells (CD11c^+ve^) from CD45.1^+^ C57BL/6 mice were pulsed with PPRV for 1hr. After washings, cells were stained to determine the presence of surface or intracellular viral antigens. Representative FACS plots show PPRV (H)^+ve^ cells for each cell type. For gating FACS plots, fluorescent minus one (FMO) parameter were used. E. Bar diagrams show cumulative data for PPRV^+ve^ cells. The experiments were repeated two more times. One way ANOVA or Student t test were done for determining the statistical significance values and are represented as following; p < 0.05 *, p < 0.01 **, p < 0.001***.

### PPRV induces lung pathologies in IFNR KO animals infected with intranasal route

PPRV causes respiratory disease in infected small ruminants and the infection spreads among animals in the herd due to their closer association, we therefore investigated its pathogenesis in mice infected via intranasal route. IFNR KO animals were infected with varying doses of PPRV and were analyzed for their survival, body weights and other physiological parameters (Fig 5). The survival analysis of animals showed a significant effect of inoculum sizes but all infected animals eventual outcome in all the animals remained same as was observed in the i.p. infected mice (Fig S2). Thus, animals infected with the lower dose (10^2^PFU) succumbed to the infection later and those infected with the higher doses (10^6^PFU) died within 6 days. For performing the cellular analysis at the tissue sites and lymphoid organs of animals, additional groups of infected animals were sacrificed when approximately 20% of their body weights were lost (Fig 5A, S3-6). WT animals were infected with a high dose (10^6^ PFU/mouse) as all i.p infected animals with the higher inoculum size survived (Fig 2B-C, data not shown). Infected WT animals reduced their body weights transiently followed by their rapid recovery until the termination of the experiments while the infected IFNR KO animals gradually reduced their body weights at both the doses (10^4^ PFU and 10^6^ PFU) of PPRV (Fig 5B). The animals were terminally anaesthetized at 6dpi for performing cellular analysis in collected BALs, lungs, brain and lymphoid organs. Approximately two fold higher infiltration level of leukocytes was observed in the BALs of WT animals as compared to the IFNR KO animals suggesting for the immune reactivity against PPRV (Fig 5C). We observed an enhanced infiltration of leukocytes in the BALs (~ 42% vs 30%) as well as lung tissues (27% vs 19%) of IFNR KO animals infected with the low (10^2^ PFU) and high (10^4^ PFU) dose of PPRV (Fig 5C and D). That the infiltrating leukocytes could be involved in protection against the virus was indicated by their inverse ratios observed in the lung tissues of two groups of PPRV infected animals (Fig 5C and D). This could suggest that an efficient viral control could be achieved before a patent lung infection is established and leukocytes are recruited. A minimal infiltration of leukocytes was evident in the brain tissues of IFNR KO and WT animals infected via intranasal route and the observed frequencies were similar in both the groups of mice (Fig 5E). We then phenotypically characterized different immune cells among the leukocyte populations in different organs. The relative abundance of macrophages (CD11b^+^F4/80^+^), neutrophils (CD11b^+^Gr1^+^), NK cells (NK1.1^+^ cells), helper (CD4^+^) and cytotoxic (CD8^+^) T cells was measured in the non-lymphoid as well as lymphoid organs of infected WT and IFNR KO mice (Fig 5F-Z). Upto a five fold reduction in the frequencies of macrophages and neutrophils were observed in the BALs, lung tissues, brain, mediastinal LNs and spleens of infected WT animals as compared to those in IFNR KO mice (Fig 5G, H, K, L, P, Q, R, W and X). The frequencies of NK cells were similar in BALs but decreased in the lung and spleen of the PPRV infected WT animals as compared to those in IFNR KO mice (Fig 5I, M and Y). The increased frequencies of both CD4^+^ and CD8^+^ T cells in BALs and lung tissues of PPRV infected WT mice in comparison to those of the IFNR KO mice were observed (Fig 5J and N). BALs and mediastinal LNs of infected IFNR KO animals injected with different doses strikingly had more frequencies of neutrophils at a lower dose as compared to those at high dose (Fig 5H and R). Such a trend was not observed in lung tissues, brain and spleen samples of the infected animals (Fig 5L, P and X). The inverse correlation with PPRV inoculum size and the recruitment of macrophages was not observed in lungs, brain and spleens of IFNR KO animals infected (Fig 5G, K O and W). Enhanced frequencies of NK cells but a reduction in the frequencies of neutrophils and macrophages were observed for both BAL and lung tissues of WT mice (Fig 5G – I and K-M). Similarly, the frequencies of both CD4^+^ and CD8^+^ T cells increased in the spleens, BALs and lung tissues of PPRV infected WT mice that efficiently controlled PPRV infection (Fig 5F, J and N). These results suggested for the anti-PPRV activity of NK cells and T cells.

**Figure 5.**
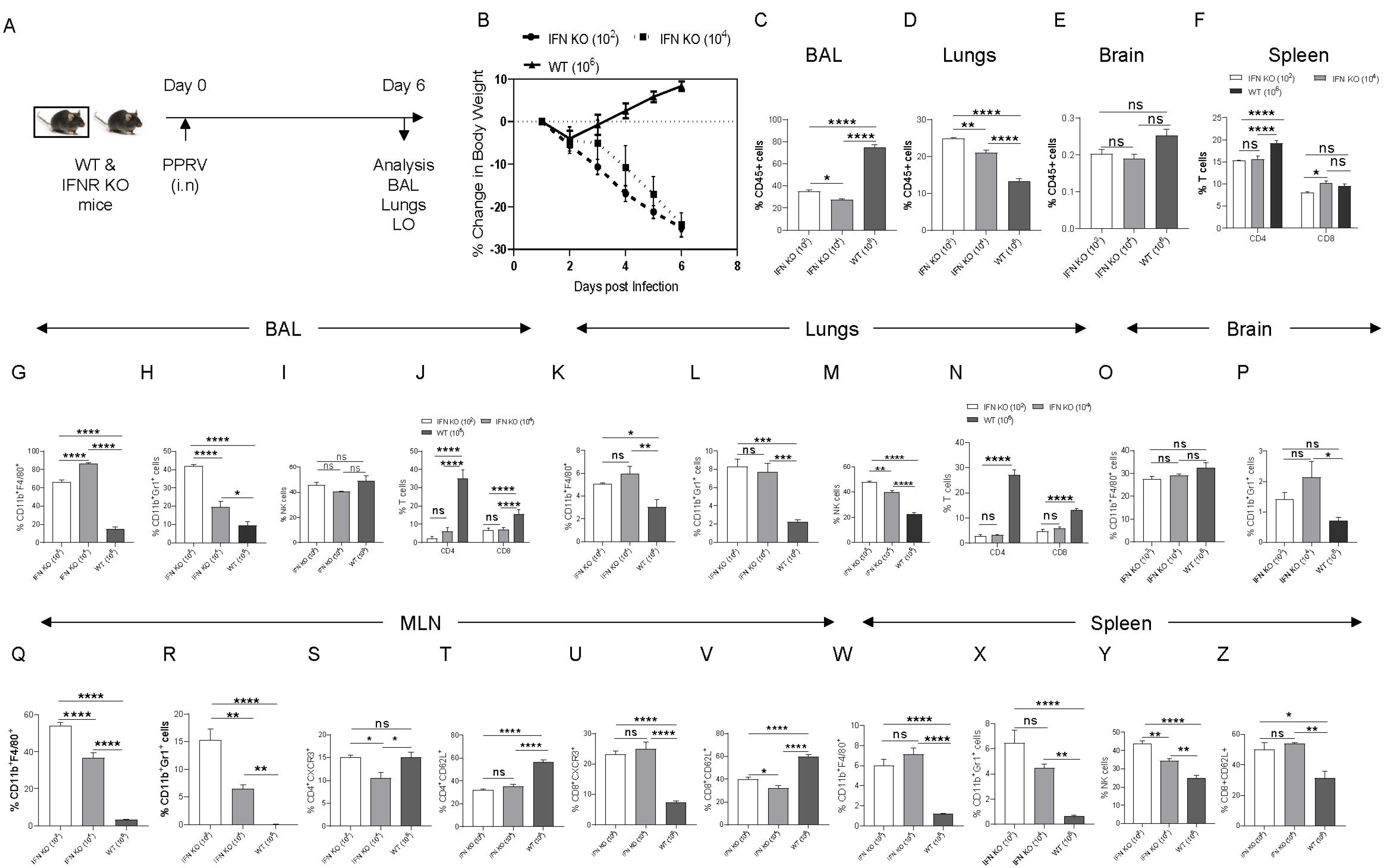
IFNR KO mice infected with PPRV develop lung pathologies and mount innate as well as adaptive immune responses. A. IFNR KO and WT C57BL/6 mice were infected with varying doses of PPRV via intranasal route and animals were analyzed for their survival, change in body weights. The level of statistical significance was determined by Gehan-Breslow-Wilcoxon test and one-way ANOVA, respectively. Terminally anaesthetized animals were analyzed for measuring the cellular infiltration in BAL, lungs and lymphoid organs at 7dpi. The FACS plots for gating strategy and representative plots from different groups of animals are shown in supplementary Fig S3-6. B. Percent change in body weights of WT and IFNR KO mice infected with PPRV is shown. C-E. The frequencies of leukocytes (CD45^+^ cells) are shown in the BAL (C), lungs (D) and brain tissues (E) of infected WT and IFNR KO mice. F. Bar diagrams show the percentage of CD4^+^ and CD8^+^ T cells in the spleens of infected animals. G-J. Frequencies of macrophages (CD11b^+^F4/80^+^) (G), neutrophils (CD11b^+^Gr1^+^) (H), NK cells (NK1.1^+^) (I) and T cells (J) are shown in the BAL of infected animals by bar diagrams. K-N. Frequencies of macrophages (CD11b^+^F4/80^+^) (K), neutrophils (CD11b^+^Gr1^+^) (L), NK cells (NK1.1^+^) (M) and T cells (N) are shown in the lungs of infected animals by bar diagrams. O-P. Frequencies of macrophages (CD11b^+^F4/80^+^) (O) and neutrophils (CD11b^+^Gr1^+^) (P) are shown in the lungs of infected animals by bar diagrams. Q-V. Frequencies of macrophages (CD11b^+^F4/80^+^) (Q), neutrophils (CD11b^+^Gr1^+^) (R) and the phenotypic characterization CD4^+^ T cells (S and T) and CD8^+^ T cells (U-V) for the indicated markers are shown in the MLN of infected animals by bar diagrams. W-Z. Frequencies of macrophages (CD11b^+^F4/80^+^) (W) and neutrophils (CD11b^+^Gr1^+^) (X) and NK cells (Y) are shown in the spleen of infected animals by bar diagrams. Z. Activation profile of splenic CD4^+^ T cells is shown by bar diagram. The experiments were repeated three times. The data was analyzed by two-way ANOVA was performed for determining the statistical significance values. The levels of significance are represented as following; p < 0.05 *, p < 0.01 **, p < 0.001 ***.

We further explored whether or not PPRV infection induces the activation of T cells. A significant proportion of CD4^+^ T cells displayed an activation profile (CD62L^lo^CXCR3^+^) in WT animals but the frequencies of such cells were up to three fold higher in the IFNR KO animals (Fig 5S and T, S5). However, the frequencies of activated CD62L^lo^CXCR3^+^ CD8^+^ T cells was ~2 fold lower in WT animals as compared to those in IFN RKO mice (Fig 5U and V, S5). Innate immune cells such as the NK cells in WT animals could help achieve an efficient viral control but such mechanisms might require the activity of IFNs as abundant replicating viral particles were present in the IFNR KO animals. Furthermore, the antigen presenting cells stimulated both CD4^+^ and CD8^+^ T cells but such cells were compromised in their function owing to their lack of type I IFN responsiveness.

Taken together, the analysis of cellular infiltration suggested for the role of neutrophils in promoting PPRV pathogenesis while NK cells and T cells playing protective roles.

### IFN responsive CD8^+^ T cells delay mortality in PPRV infected IFNR KO mice

We established the susceptibility of IFNR KO mice to PPRV, the potential spread of PPRV by infected innate immune cells and expansion of innate immune cells as well as T cells in PPRV infected WT mice that controlled the virus well (Fig S6B). We then explored whether WT T cells could either protect infected IFNR KO animals or reduce the severity of PPRV infection. FACS sorted CD4^+^ and CD8^+^ T cells from WT mice were transferred into IFNR KO animals. Such cells were allowed to expand for 40 days in recipients, which were then infected with PPRV (Fig S7A). In comparison to infected controls that gradually lost their body weight, CD8^+^ T cell recipients showed a significantly reduced body weights until their termination at 6dpi (Fig S7B). WT CD4^+^ T cell recipient mice and those received bone marrow cells did not shown alteration in the rates or the kinetics of body weight loss (Fig S7B, data not shown). The activation status of CD8^+^ and CD4^+^ T cells of control and T cell recipient mice revealed more frequencies of CD4^+^ and CD8^+^ T cells expressing high levels of CD44 when WT CD8^+^ T cells were transferred (Fig S7C-F, data not shown). These results suggest anti-viral activity of WT CD8^+^ T cells.

We then measured whether or not previously PPRV-primed WT CD8^+^ T cells confer protection to the PPRV infected IFNR KO mice. 5×10^6^ of FACS sorted WT CD8^+^ T cells from naïve or the mice previously infected with PPRV were transferred before infecting IFNR KO mice with PPRV (Fig 6A). The survival analysis showed a significant advantage conferred to the infected IFNR KO mice by transferred naïve or primed CD8^+^ T cells in delaying mortality (Fig 6B). Such effects occurred in a dose dependent manner with animals receiving five fold lower CD8^+^ T cells succumbed to the infection early (Fig S7 and data not shown). Separate groups of PPRV infected CD8^+^ T cell recipients were measured for a change in their body weights and cellular analysis on 7dpi (Fig S8 and S9). CD8^+^ T cell recipients showed significantly reduced body weights by 6dpi (Fig 6C). Cellular analysis showed that approximately 50% of CD45^+^ cells were present in the BAL of PPRV infected animals and the frequencies decreased to 30% in CD8^+^ T cells recipients (Fig 6D). Similarly in the lung tissues of infected animals the frequencies of CD45^+^ cells reduced from 25% to 15% in CD8^+^ T cells recipients (Fig 6E). However, such effects were prominently observed in the group receiving CD8^+^ T cells from previously PPRV-infected animals. Neutrophils levels increased in the BALs of CD8^+^ T cell recipients IFNR KO mice by ~1.5 fold but the macrophages and NK cells decreased by ~ 10% (Fig 6F(a)-F(c)). In the lungs tissues, no significant differences in the cellular infiltrations were observed but for an increase in macrophages in CD8^+^ T cell recipients (Fig 6G). The cellular analysis in the MLN and spleen samples of infected animals revealed a reduction in the frequencies of innate immune cells such as neutrophils and macrophages but more CD4^+^ and CD8^+^ T cells were observed in WT CD8^+^ T cell recipient mice (Fig 6J and H). Phenotypic characterization of CD4^+^ and CD8^+^ T cells in control and WT CD8^+^ T cell recipient PPRV infected IFNR KO mice showed a reduced expression of the activation molecule CXCR3 and as well as a lymph node retention molecule CD62L (Fig 6I). However in the spleen of WT CD8^+^ T cell recipient mice the expression of CXCR3 was increased but that of CD62L reduced by both CD4^+^ and CD8^+^ T cells (Fig 6K). These results might mean suggest for the retention of CD8^+^ T cells in the LNs of infected animals probably due to more chemokines build up and a less efficient gradient generation for such chemokines to facilitate their exit and viral control at the infection sites (25). A detailed analysis is currently underway.

**Figure 6.**
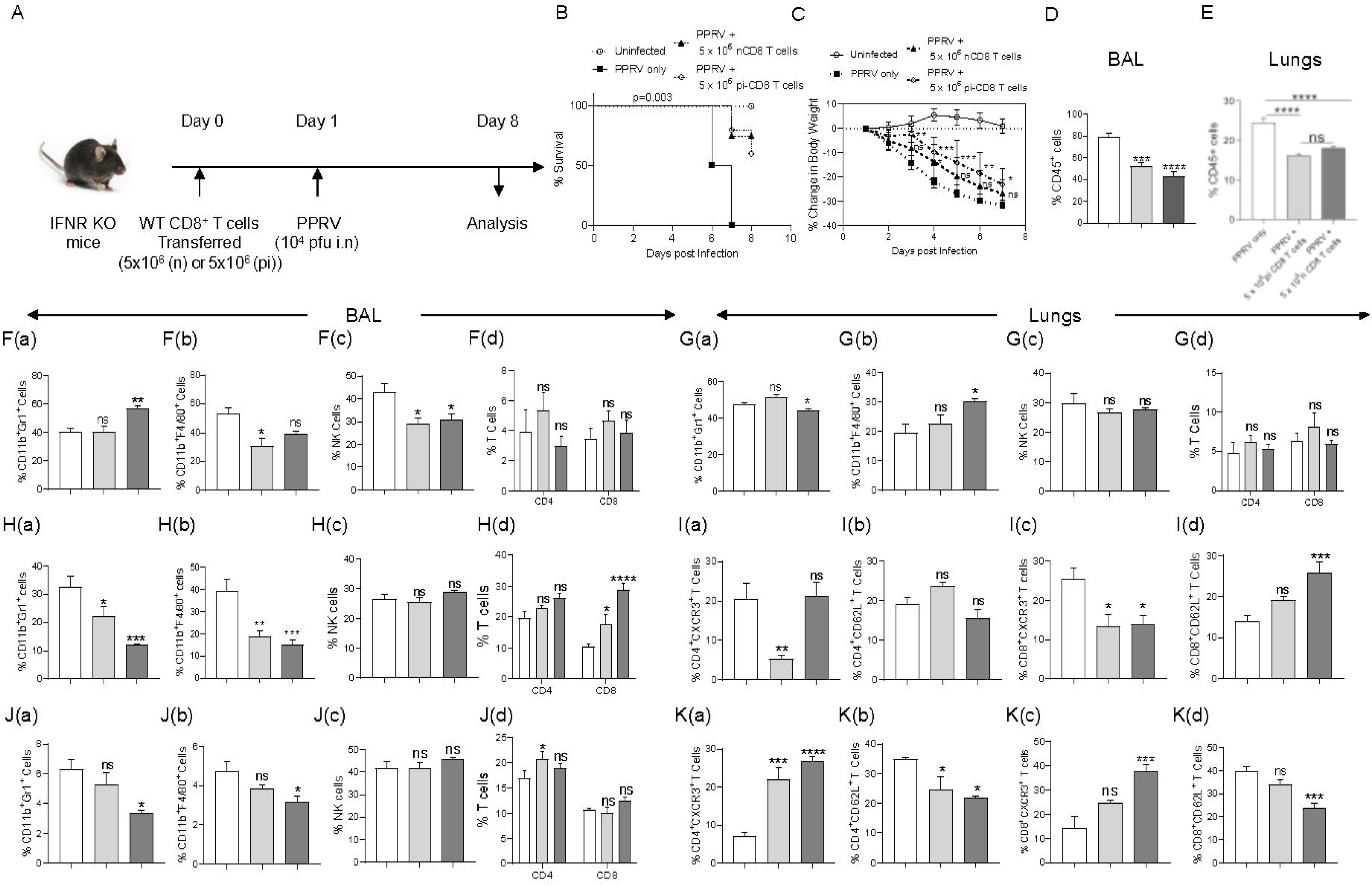
WT CD8^+^ T cells delay mortality in PPRV infected IFNR KO mice. A schematic of the experiments is shown. 5×10^6^ of CD8^+^ T cells from naïve or previously PPRV infected WT mice were transferred into IFNR KO mice. Next day, the recipient animals were infected with PPRV (10^4^PFU). The disease progression, survival and cellular analysis of immune cells were performed. B. The survival in different groups of animals is shown. The results were analyzed by Gehan-Breslow-Wilcoxon test. C. Percent change in body weight of mice from each group is shown. The level of statistical significance was determined by Two-way ANOVA. D-E. The frequencies of leukocytes (CD45^+^ cells) in BAL (D) and lungs (E) of infected mice are shown by bar diagrams. F-G. The frequencies of macrophages (CD11b^+^F4/80^+^), neutrophils (CD11b^+^Gr1^+^), NK cells (NK1.1^+^) and T cells (both CD4^+^ and CD8^+^ T cells) are shown in the BAL (F) and lungs (G) of infected animals by bar diagrams. H. The frequencies of macrophages (CD11b^+^F4/80^+^), neutrophils (CD11b^+^Gr1^+^), NK cells (NK1.1^+^), T cells (CD4^+^ and CD8^+^ T cells) are shown in the MLNs of infected animals by bar diagrams. I. The phenotypic characterization CD4^+^ T cells and CD8^+^ T cells for the indicated markers are performed and the percent positive cells frequencies for the indicated markers are shown for MLN of infected animals by bar diagrams. J. The frequencies of macrophages (CD11b^+^F4/80^+^), neutrophils (CD11b^+^Gr1^+^), NK cells (NK1.1^+^), T cells (CD4^+^ and CD8^+^ T cells) are shown in the MLNs of infected animals by bar diagrams. K. The phenotypic characterization CD4^+^ T cells and CD8^+^ T cells for the indicated markers were performed and the frequencies of percent positive cells for the indicated markers are shown for spleen of infected animals by bar diagrams. The experiments were performed three times. The data was analysed by two-way ANOVA for determining the statistical significance values and are represented as following; p < 0.05 *, p < 0.01 **, p < 0.001 ***.

We observed an aberrant activation profile of T cells in PPRV infected mice, but a crucial role of functionally competent CD8^+^ T cells was evident in mitigating PPRV pathogenesis.

### Identification of immunogenic CD8^+^ T cell epitopes of PPRV *in silico*

We identified immunogenic epitopes of PPRV that could induce specific CD8^+^ T cells response in mice. All the structural proteins (heamagglutinin, matrix, nucleocapsid and fusion proteins) of PPRV were analyzed for predicting H-2K^b^ restricted epitopes using IEDB database. We focused our analysis on the structural proteins because such proteins are critical for the viral assembly and its envelope formation (24). Peptides with low percentile ranks and the IC_50_ value of <200nM were selected (Table S1). Out of the list generated, top 12 best ranking peptides (three from each protein) were chosen for synthesis and further analysis (Fig 7A). The peptides included in analysis were IVVRRTAGV, VAFNILVTL, FMYLFLLGV (matrix protein), FSAGAYPLL, ASFILTIKF, SSITTRSRL (Nucleocapsid protein), VILDRERLV, IEHIFESPL, IGLVRDFGL (hemagglutinin protein) and (AILTFLFLL, VAILTFLFL, SGGDFLAIL (fusion protein). The peptides were subjected to *in silico* analysis. Molecular docking is one of the most frequently used methods to predict the conformation of small-molecule ligands. Docking of selected peptides from different proteins of PPRV against H-2K^b^ was performed and the models were analyzed using Chimera tool. The energy parameters of the best fitting structures among PPRV peptides were determined (Table S2 and S3). The best docking results were provided by PPRV peptides FMYLFLLGV and FSAGAYPLL with H-2K^b^. FMYLFLLGV peptide even yielded better docking scores then the reference peptide FAPGNYPAL (SEV-9) of Sendai E virus (Fig 7B and D and Table S2). Further analyses revealed phenylalanine residues at position 1 and 5 in FMYLFLLGV peptide as the probable anchors. Similarly, for FSAGAYPLL peptide phenylalanine at position 1 and proline at position 7 were predicted to be the most probable anchors. Docking of peptides was also performed with class MHC molecule of goat (CLA1) (Fig 7C and F, S10 and Table S3). CLA-I and H-2K^b^ superposed near perfectly at an RMSD value of 1.5A (Fig 7E). The predicted peptides showed a similar trend of docking with CLA-1 molecule. The anchor residues as well as the docking scores of FMYLFLLGV and FSAGAYPLL peptides scored better in these analyses (Fig 7E-F). Interestingly, all the epitopes displayed better docking with CLA-1 than with H-2K^b^ indicating their immunogenicity in generating anti-viral response against PPRV infection in the natural host, goats (Fig 7F). A total of 12 peptides were predicted that could potentially be immunogenic in mice as well as in the natural host of PPRV goats.

**Figure 7.**
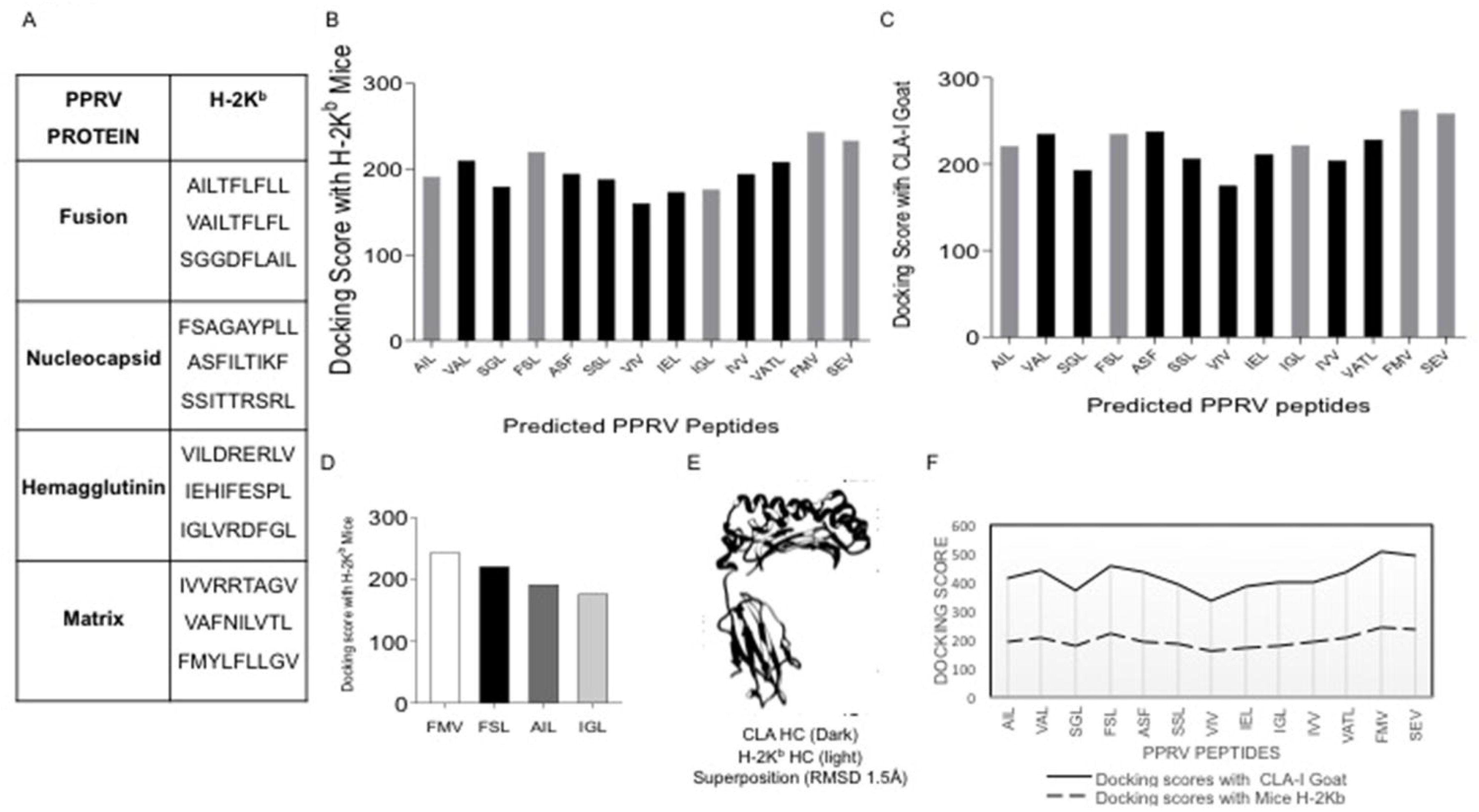
Molecular docking reveals docking of predicted peptides binding with H-2K^b^ and CLA-I. A. High scoring peptides and their origin of PPRV protein are tabulated. B. The docking scores of each peptide with H-2K^b^ are shown. C. The docking scores of each peptide with goat class I MHC (CLA-1) are shown by bar diagrams. D. Selected peptides for further evaluation in *ex vivo* and *in vivo* assays with their predicted molecular docking scores are shown. E. H-2K^b^ and CLA-1 were superimposed at RMSD value of 1.5A. F. A comparative analysis of docking scores of H-2K^b^ and CLA-I for the predicited PPRV peptides is shown.

### Assessing class I MHC stabilization potential of predicted PPRV epitopes using acellular and cellular assays

We tested all top performing peptides for their class I MHC stabilization potential using acellular and cellular assays. SIINFEKL, an Ova derived peptide with known immunogenicity for H-2K^b^, was used as a positive control. In order to measure the MHC stabilizing potential of predicted epitopes, we performed ELISA. The heterotrimeric complex consisting of a photocleavable ligand, β2 microglobulin and H-2K^b^ was immobilized to solid phase by plate-coated streptavidin. UV displaced conditional ligand and its replacement with the testing peptide yields a positive reaction that can be detected by anti-β2 microglobulin antibody (16). Fold change values as compared to those obtained for SIINFEKL peptide for each of the PPRV derivative peptides are shown in Fig 8A. Accordingly, FSAGAYPLL, IGLVRDFGL, AILTFLFLL, SSITTRSRL, IVVRRTAGV, IEHIFESPL, VAFNILVTL, and VILDRERLV displayed higher values as compared to other peptides (Fig 8A).

**Figure 8.**
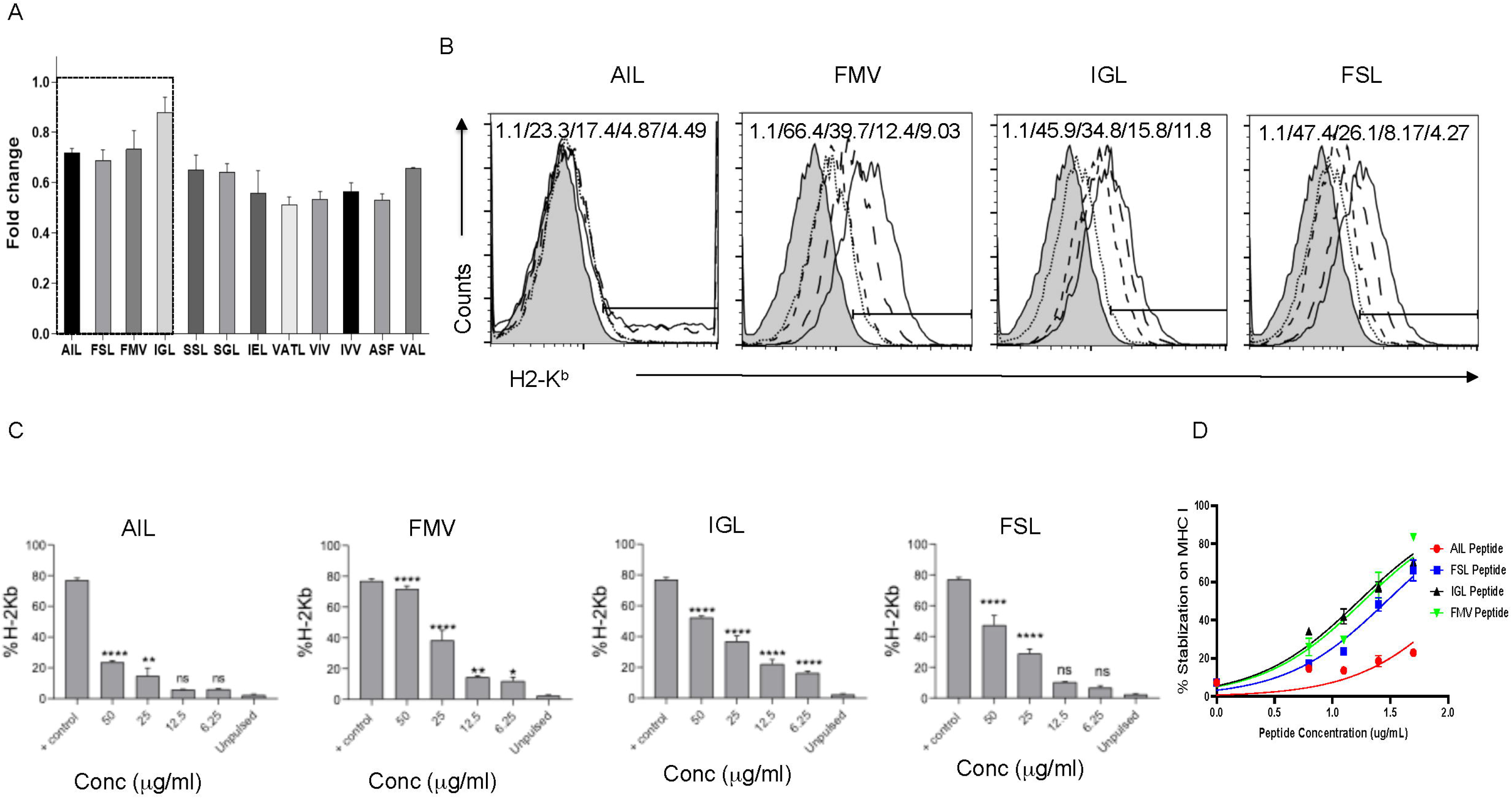
Immunogenicity of predicted peptides of PPRV was determined by *in vitro* assays. A. An ELISA was performed for measuring the stabilization of H-2K^b^ monomers by synthetic peptides predicted to be immunogenic. The bar diagrams show folds change values for H-2K^b^ positive cells for the respective peptides as compared to the control peptide. B. Representative overlay histograms for different concentrations (50μg/ml, 25μg/ml, 12.5μg/ml and 6.25μg/ml) of indicated peptides show their ability to stabilize surface MHC molecules. The values represent the percent positive cells at different concentrations of peptides. C. Bar diagrams show percent H-2K^b^ positive cells for different concentrations of the indicated peptides. D. Line graph shows log EC_50_ values for different peptides for stabilizing H-2K^b^ in pulsed RMA/s cells. Student t test were performed for determining the statistical significance values and are represented as following; p < 0.05 *, p < 0.01 **, p < 0.001 ***.

The peptides were also analysed for their class I MHC stabilizing activities using transporter associated with antigen processing and presentation (TAP) deficient RMA/S cells. Such cells express fewer molecules of class I MHCs on surface but the exogenously added immunogenic peptides help stabilize their expression (26). Different concentrations PPRV peptides were added to serum starved RMA/S cells (Fig 8B-D). Out of 12 peptides, four peptides FMYLFLLGV (matrix protein), FSAGAYPLL (nucleocapsid protein), IGLVRDFGL (hemagglutinin protein) and AILTFLFLL (fusion protein) showed higher affinity for H-2K^b^ molecule and induced more cells to express H-2K^b^ (Fig 8B-D). The results were dependent on the concentrations of the peptides used (Fig 8B-D). Similar results were obtained when the mean fluorescence intensity (MFI) values were measured for the expression of H-2K^b^ by each peptide (data not shown). Log EC_50_ values for AILTFLFLL, FSAGAYPLL, IGLVRDFGL and FMYLFLLGV peptides were 2.098, 1.469, 1.228 and 1.268 μg/ml, respectively (Fig 8D).

### Immunogenicity of PPRV peptides in PPRV infected or immunized mice

We measured response of CD8^+^ T cells in PPRV infected and immunized mice. As C57BL/6 mice were refractory to PPRV infection, we used a high dose of PPRV (5×10^6^ PFU) PPRV for infection. After 7 days, a boosting dose was given and the analysis was done in blood samples three days later by measuring the numbers and frequencies of class I MHC tetramer (H-2K^b^-p(PPRV)-tetramer). Our results showed significantly more numbers of antigen-specific CD8^+^ T cells against AILTFLFLL, FSAGAYPLL and IGLVRDFGL peptides of PPRV as compared to those induced against FMYLFLLGV (Fig 9A-C). Tetramer positive CD8^+^ T cell count was also high for FSAGAYPLL, IGLVRDFGL peptides in animals immunized with the cocktail of four peptides (Fig. 9D-E). The functionality of PPRV specific CD8^+^ T is currently being investigated.

**Figure 9.**
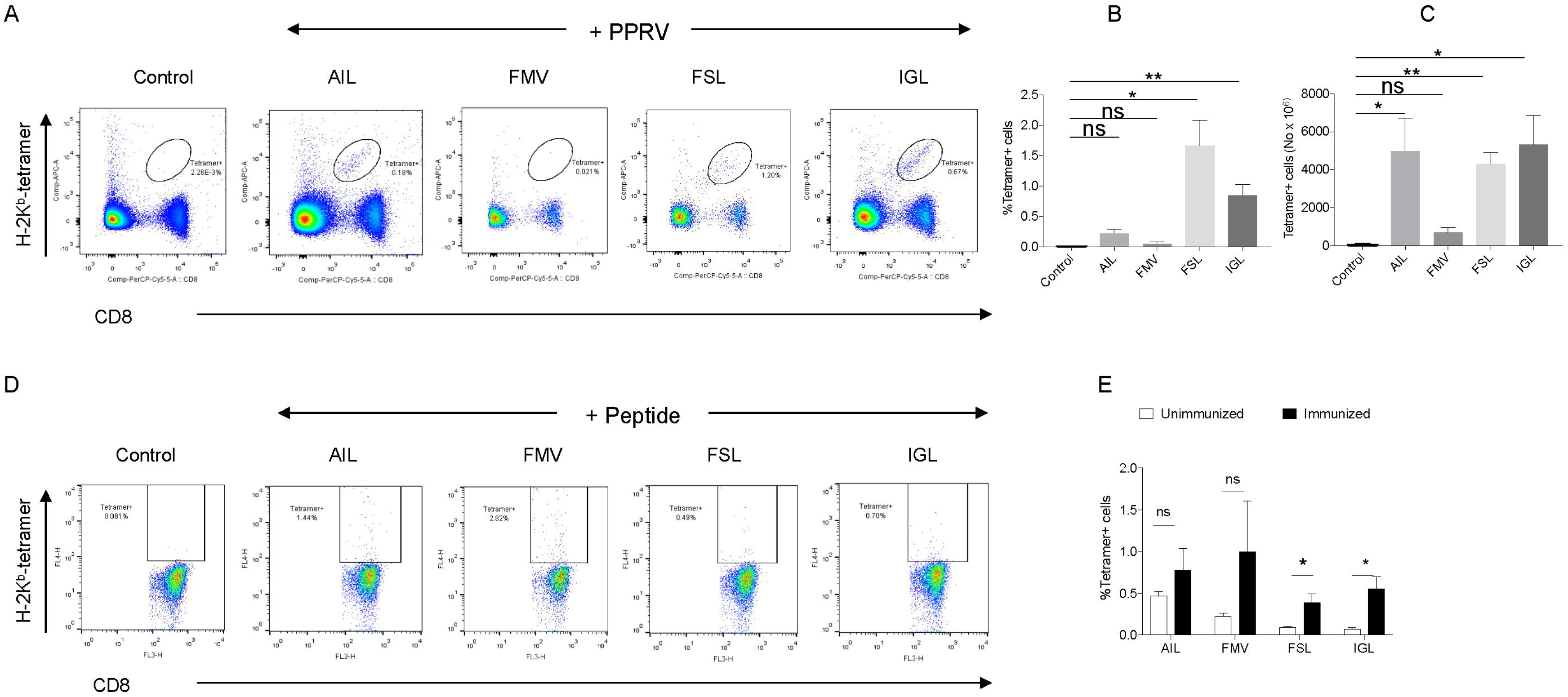
PPRV infected and PPRV-peptide immunized WT C57BL/6 mice expand virus-specific CD8^+^ T cells. A-C. WT C57BL/6 mice (n=5) were i.p. infected with 5×10^6^ PFU of PPRV at an interval of 7 days. The analysis of the expanded cells was performed three days later by measuring the frequencies PPRV-specific CD8^+^ T cells by staining with arrays of tetramers. A. Representative FACS plots shown the frequencies of class I MHC tetramer positive cells for the indicated peptides. B. Frequencies of PPRV-specific CD8^+^ T cells are shown by bar diagrams. C. The number of cells/million of PBMCs is shown by bar diagrams. D-E. C57BL/6 mice (n=4) were immunized either with the cocktail of peptide (AILTFLFLL, FMYLFLLGV, FSAGAYPLL and IGLVRDFGL) each with 5μg/mouse in complete Freund’s adjuvant via subcutaneous route. After two weeks, a second injection of the same concentration was administered as an emulsion with incomplete Freund’s adjuvant. Three days later the frequencies of peptide specific CD8^+^ T cells were measured by tetramer staining of PBMCs. D. Representative FACS plots show the frequencies of PPRV-specific CD8^+^ T cells. E. The number of cells/million of PBMCs is shown by bar diagrams. The experiments were repeated two times and the statistical significance was measured by Student t test. p < 0.05 *, p < 0.01 **.

Our results suggest an induction of CD8^+^ T cell response against predicted PPRV epitopes. Therefore, our results also suggest that immunization with peptides could represent a potential subunit vaccination approach for PPRV to investigate properties of CD8^+^ T cells during PPRV infection in mice.

## Discussion

With ramped up efforts to eradicate PPRV by intensive vaccination programs, it has become imperative to develop an accessible laboratory animal model to better understand its pathogenesis and more importantly the immune correlates of protection against the virus. Such investigation could enhance prospects of devising an alternative vaccine strategies should a need arise. We undertook this study to develop a laboratory mouse model to elucidate PPRV pathogenesis, the role of innate immune cells and cytotoxic T cells in its control. We demonstrated the susceptibility of IFNR KO mice to PPRV infection. The infected animals succumbed to PPRV infection irrespective of the dose of inoculum and route of infection. The replicating viruses as well as the derivative antigens were present abundantly in most of the critical organs of infected mice. Neutrophils and macrophages likely served as the Trojan horse to transport virus to the CNS to cause encephalitis while CD8^+^ T cells in addition to NK cells were protective in PPRV infected mice. We also discovered immunogenic epitopes of PPRV and enumerated virus-specific CD8^+^ T cells in infected and immunized mice C57BL/6 mice using an array of MHC tetramers. Our results showed the infectivity of adult mice that can serve as a laboratory animal model for investigating PPRV pathogenesis and further decipher protective role of CD8^+^ T cells.

PPRV, a member of morbilivirus genus of paramyxoviridae family, incurs significant losses to animal husbandry sector in endemic areas (27). Therefore, intensive vaccination programs are being adopted in many countries to eradicate the virus. A laboratory mouse model would be useful not only to better understand the contribution of cellular and molecular mediators in the viral of pathogenesis but also to test the efficacy of anti-virals. One such class of host-derived molecules includes type I IFNs that have potent anti-viral effects. We observed distinct expression pattern of type I IFNs induced by PPRV stimulated innate immune cells depending on the dose of infecting virus, and the intrinsic properties of responding cells (RAW macrophages versus bone marrow derived primary macrophages). These results could suggest for a diversification in the function of type I IFNs. Such a phenomenon is well documented for the activity of type I and type III IFNs that was shown to be largely attributed to the expression pattern of their cognate receptors (28). While the receptors for type I IFNs (IFNAR1 and IFNAR2) are ubiquitously present on most cells, type III IFNRs (IFNL complex) are predominantly present in epithelial cells and only a subset of innate immune cells such as neutrophils (27, 29). A dichotomy is also known with the activity of type I IFNs i.e., IFNα and IFNβ(30). Furthermore, a specific inhibition of IFNβsignaling by antibodies converted the course of a persistent viral infection with LCMV clone 13 into an infection that could be efficiently controlled in the acute phase (31). Therefore, further analysis to decipher the relative roles of different species of type I IFNs in the protection against PPRV would be valuable.

In WT animals the activity of NK cells might control PPRV infection sufficiently and CD4^+^ and CD8^+^ T cells playing a subsidiary antiviral role (Fig 5). We observed efficient infiltration of NK cells in the BAL and lung tissues but the animals eventually succumbed to the infection (Fig 5I and M). These results suggested a critical role of IFN signaling in the NK cells mediated control of PPRV and the killing activity of such cells alone might not suffice to achieve viral control. The expansion and the migration of T cells particularly of CD4^+^ T cells in PPRV infected WT mice in comparisons to those in the IFNR KO mice could suggest for a critical role played by IFN signaling in the efficient activation or the migration of such cells which can then help efficient priming of CD8^+^ T cells. That the CD4^+^ T cell responses precede CD8^+^ T cells was shown earlier (32). Earlier reports also suggested that an inefficient JAK/STAT signaling in CD8^+^ T cells that occur following type I IFN ligation with their cognate receptors enhances the propensity of such cells to undergo apoptosis by host factors such as the glucocorticoids. Multiple studies have shown that microbial infections can actively engage hypothalamic pituitary adrenal (HPA) axis to induce glucocorticoids able to induce the apoptosis of CD8^+^ T cells (17). Whether or not PPR infection predominantly activate HPA axis is not known currently. That CD8^+^ T cells are involved in anti-PPRV defense mechanisms was shown by the adoptive transfer of WT CD8^+^ T cells, isolated from previously infected mice that resolved the infection (Fig 6).

With limited data available for PPRV cell and tissue tropism, its replication was demonstrated in the lymphoid organs (6). The known receptor for PPRV are SLAM family proteins (33). Our results demonstrated that the cells of innate immune origin expanded upon PPRV infection in IFNR KO mice were susceptible to the viral infection. Surprisingly innate immune cells such as neutrophils and macrophages were potentially able to transfer virus to distal locations (Fig 3 and 4). The innate immune cells express SLAM receptors abundantly and therefore their susceptibility to PPRV infection could involve these receptors (33). The identification of a particular receptor in the susceptibility of mice has not been explored and constitutes part of our ongoing investigations. It would be interesting to explore whether PPRV replicates in the innate immune cells of sheep and goats. If indeed such cells exhibit susceptibility to PPRV, it might necessitate revisiting vaccination strategies to help achieve a complete viral eradication. Neurovirulence and neuropathology induced by PPRV are recently reported in goat kids and newly born BALB/c mice as well as CD1d knockout mice, the latter induce inefficient NK cells responses (22, 23). However the susceptibility of IFNR KO mice to PPRV was not shown earlier. Our observations could be more relevant for some herds that might have mutations in one or more components of signaling pathways involving type I IFNs. Animals and humans with signaling defects in pathways leading to type I IFN production are exceedingly susceptible to viral infections (21, 34). We used a vaccine strain of PPRV that induced a hundred percent mortality in IFNR KO mice and no apparent disease in WT C57BL/6 mice. Whether or not the virulent strain of PPRV can induce the disease and potent CTL response in WT mice is currently being investigated in our laboratory.

An initial encounter of host with viruses elicit type I IFN response, but for a long term protection the optimal activity of CD8^+^ T cells is crucial in virus clearance (17, 35). This necessitates identifying class I epitopes that can be used to quantify and assess functionality of such cells (11). Moreover, designing a vaccine against an intracellular pathogen also requires information about immunogenic epitopes that can also serve as subunit vaccine candidates. Screening of peptides of an antigen is best done by *in silico* analysis as invariably a large number of linear amino acid sequences need to be probed. The utility of class I MHC tetramers in staining antigen-specific CD8^+^ T cells and discovering epitopes in a throughput manner is unmatched but has not been adequately put to use particularly for animal pathogens (18, 36). The prediction of peptide is necessary for diagnosis, formulating vaccines as well as for analyzing the functionality of cells (11, 18, 37). We discovered at least four immunogenic epitopes from structural proteins of PPRV using *in silico, ex vivo* and *in vivo* approaches. That the immunization of mice with a cocktail of peptides induced PPRV specific CD8^+^ T cells constitute first such example for PPRV. Furthermore such an approach provides impetus to subunit vaccine development. Our ongoing investigations would help decipher pathways in virus specific CD8^+^ T cells that could help provide protection during acute as well as memory response.

## Notes

**Conflict of financial interest:** The authors declare no conflict of financial interest.

The study was supported by extramural grant from National Agriculture Science Fund (NASF/ABA-6021/2017-18).

### Competing Interest Statement

The authors have declared no competing interest.

