## Supplementary Figures for "Modeling PPRV pathogenesis in mice to assess the contribution of innate cells and anti-viral T cells"

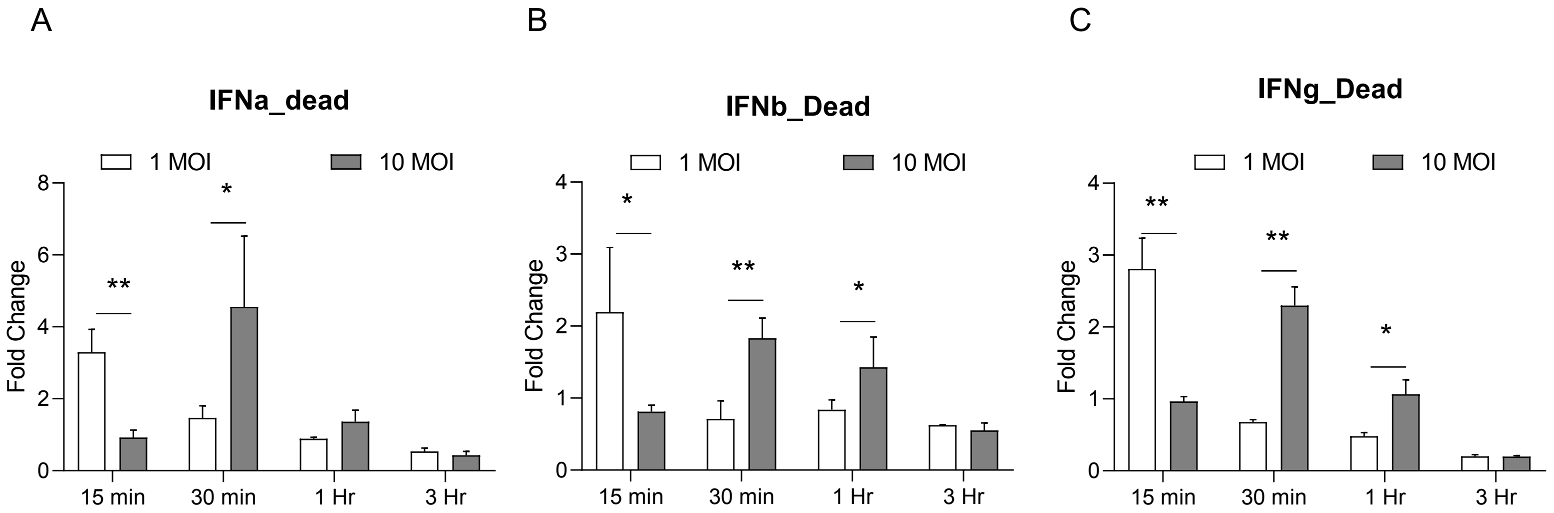


Figure S1. Heat inactivated PPRV was used for pulsing RAW macrophages and the IFN response was measured at mRNA levels by qRT PCR

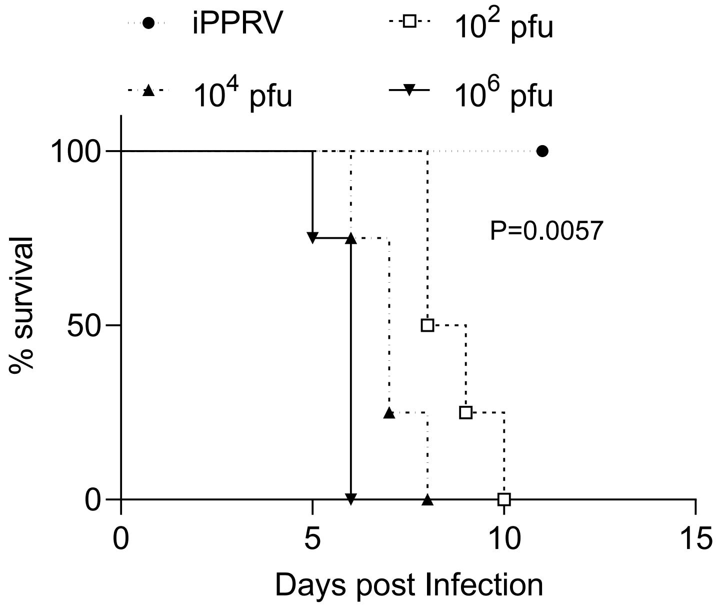


Figure S2. Survival analysis in IFNR KO mice i.p infected with varying doses of PPRV

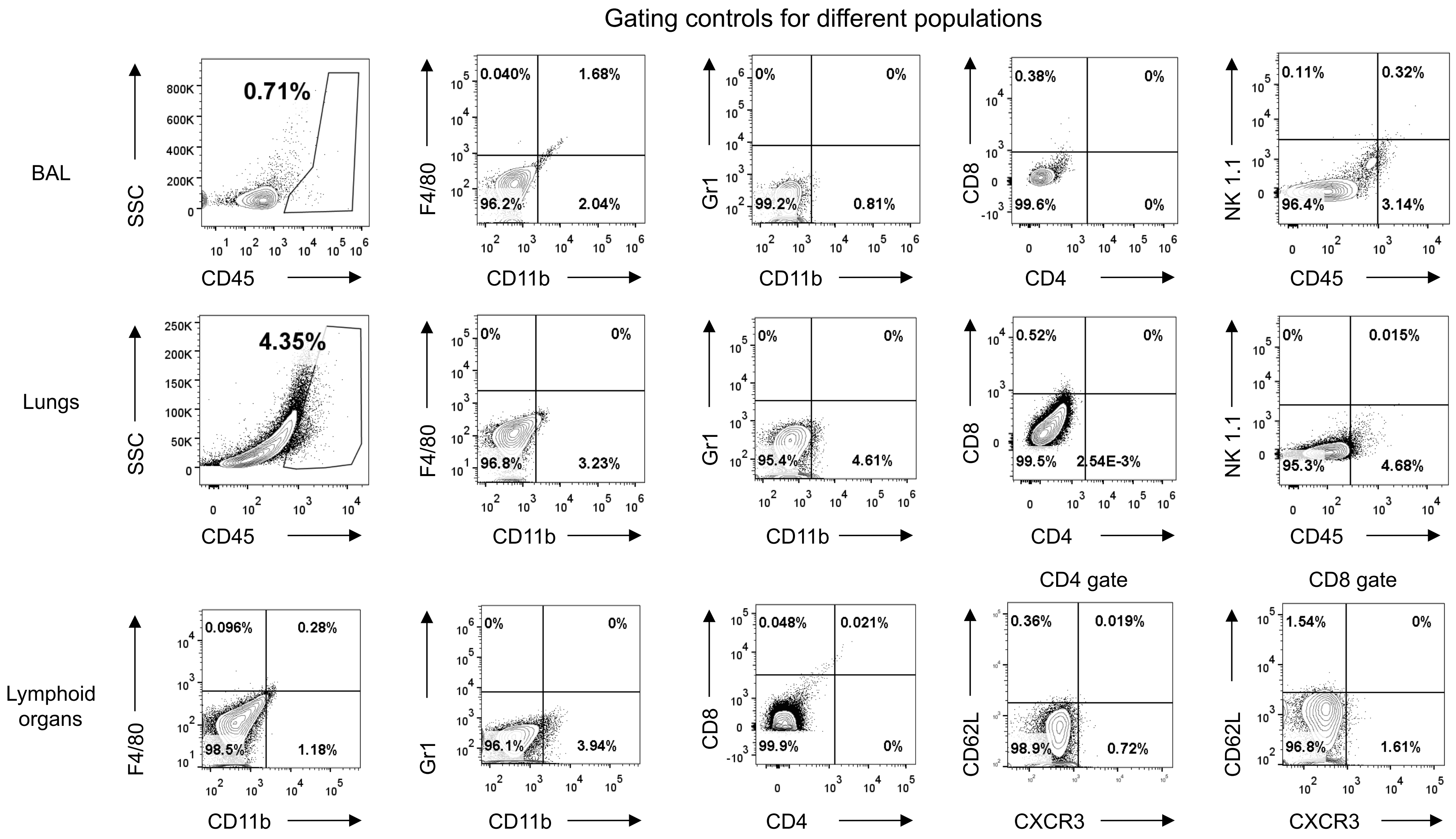


Figure S3. Gating strategy for analyzing different immune cells in lymphoid and non-lymphoid organs. Indicated markers are shown for the respective FACS plots.

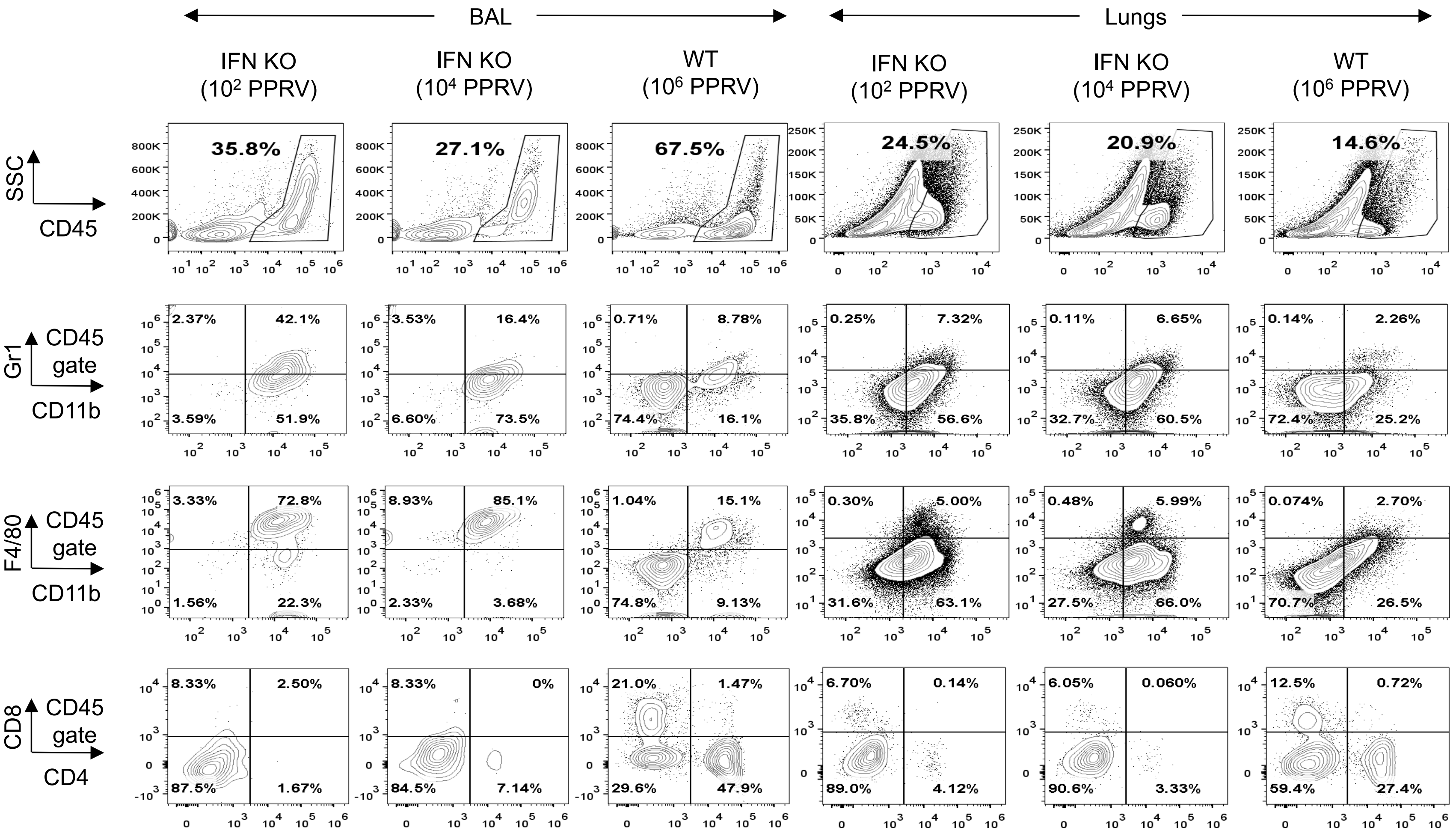


Figure S4. Representative FACS plots show the cellular distribution in bronchoalveolar lavage (BAL) and lung tissues of PPRV infected IFNR KO mice. The animals were infected with the indicated doses of PPRV via intranasal route and scarified on 6dpi and the cellular analyses were performed in BAL and lung tissues. Indicated markers are shown in the respective FACS plots.

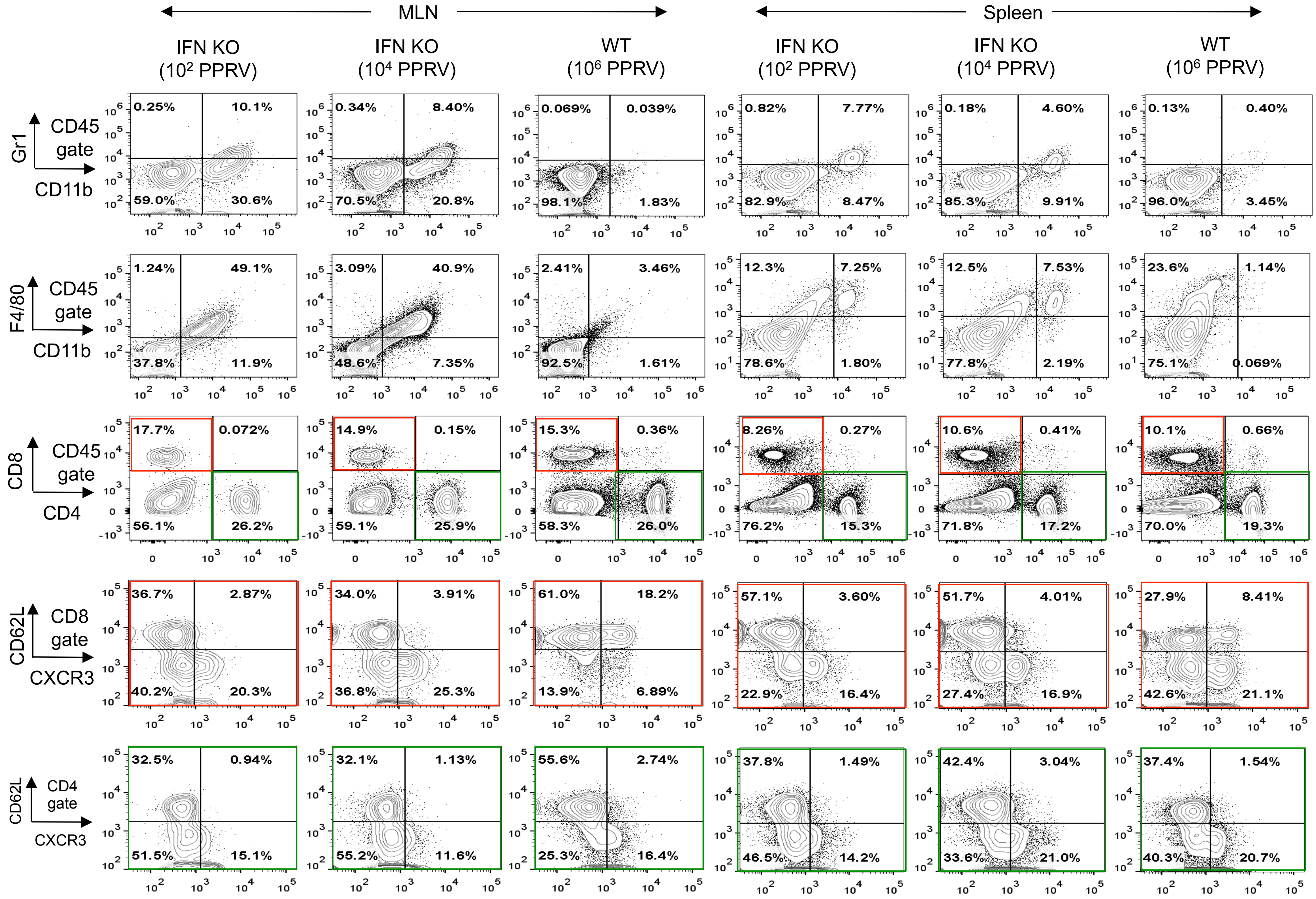


Figure S5. Representative FACS plots show the cellular distribution in mediastinal lymph nodes (MLN) and spleen of PPRV infected IFNR KO mice. The animals were infected with the indicated doses of PPRV via intranasal route and scarified on 6dpi and the cellular analyses were performed in single cell suspension of MLN and spleen. Indicated markers are shown in the respective FACS plots.

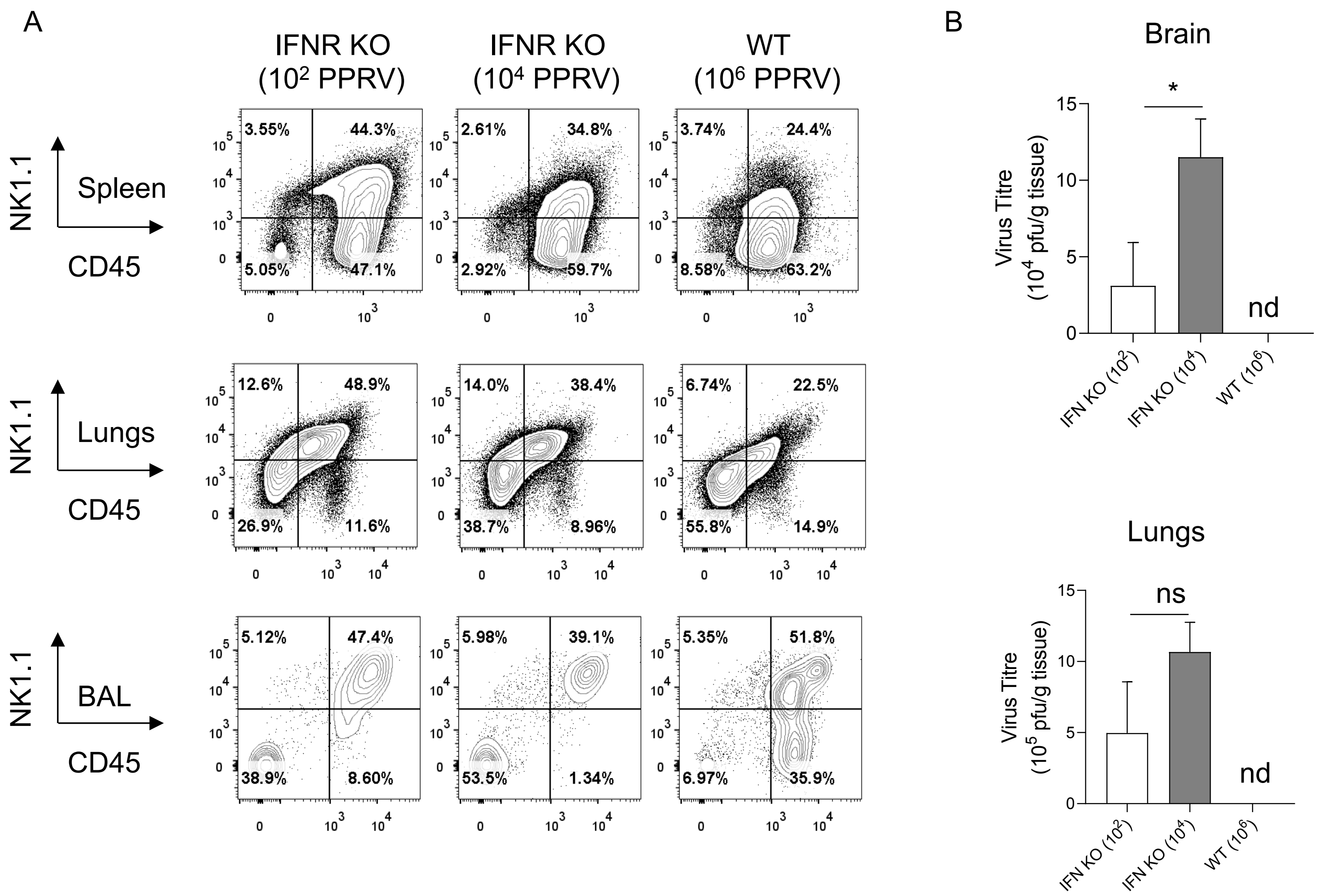


Figure S6. Representative FACS plots show the distribution of NK cells in indicated organs of PPRV infected IFNR KO mice. The animals were infected with the indicated doses of PPRV via intranasal route and scarified on 6dpi. A. The cellular analyses were performed in single cell suspension of spleen, lungs and BAL. Indicated markers are shown in the respective FACS plots. B. The viral titers measured by plaque assays are shown by bar diagrams. nd= not detectable levels.

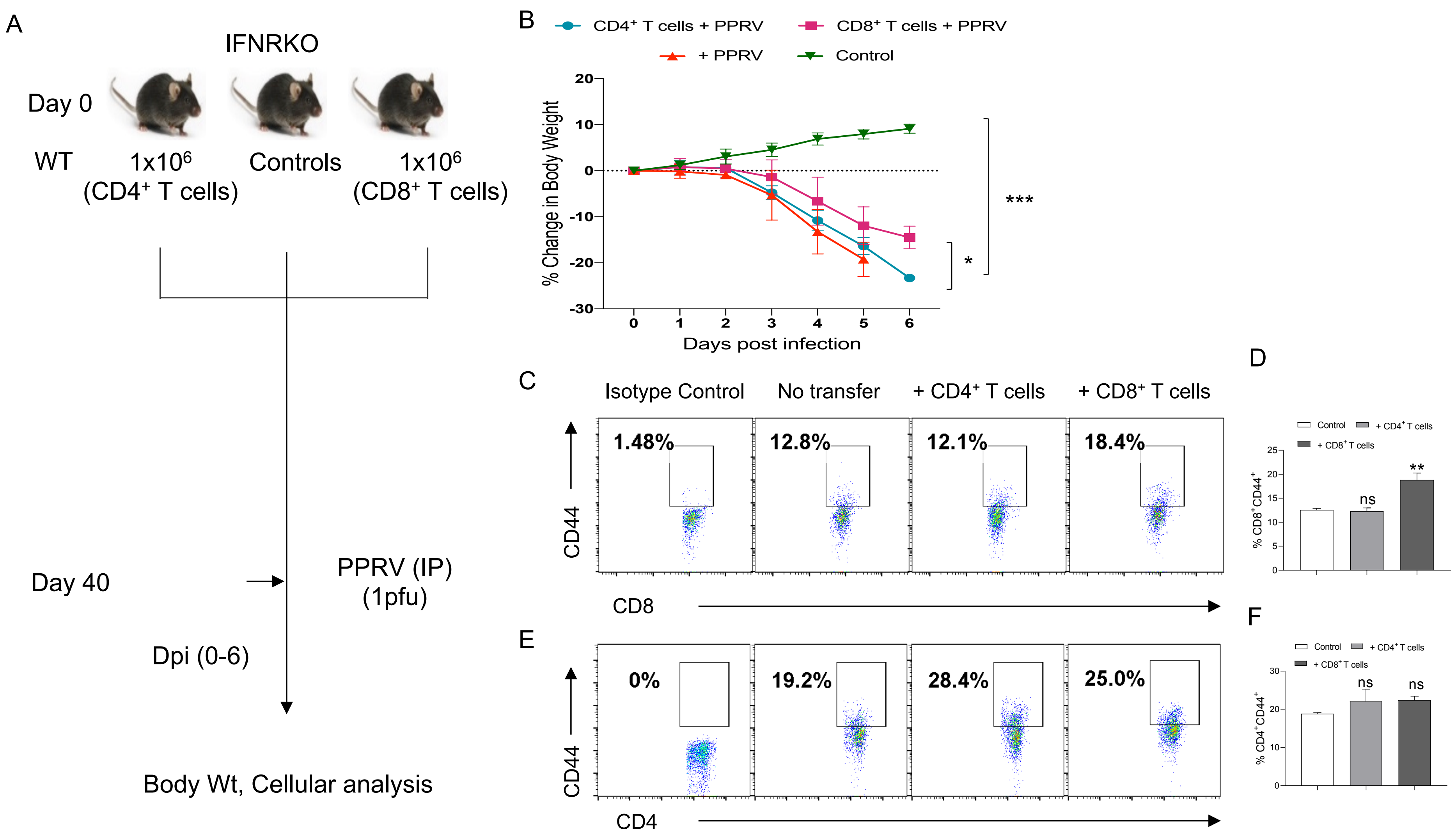


**Figure S7. WT CD8⁺ T cells delay morbidity in PPRV infected IFNR KO mice. A.** A schematic of the experiments is shown. $1x10^6$ $CD4^+$ or $CD8^+$ T cells from WT mice ($CD45.1^+$) into IFNR KO mice ($CD45.2^+$). After 40 days, animals were i.p infected with PPRV. The disease progression and the activation profile of $CD4^+$ and $CD8^+$ T cells were measured. B. Percent change in body weight of mice from each group is shown. The level of statistical significance was determined by one-way ANOVA test. C. Representative FACS plots show the activation of $CD8^+$ (C) and $CD4^+$ T cells (D) recovered from the spleen samples in different groups. D. In each group four animals were used. Cumulative data on the activation profile of $CD4^+$ and $CD8^+$ T cells is shown by bar diagrams. Student t test was used for analysis. The experiments were repeated two more times with essentially similar results. $p < 0.05$ *, $p < 0.01$ **, $p < 0.001$***.

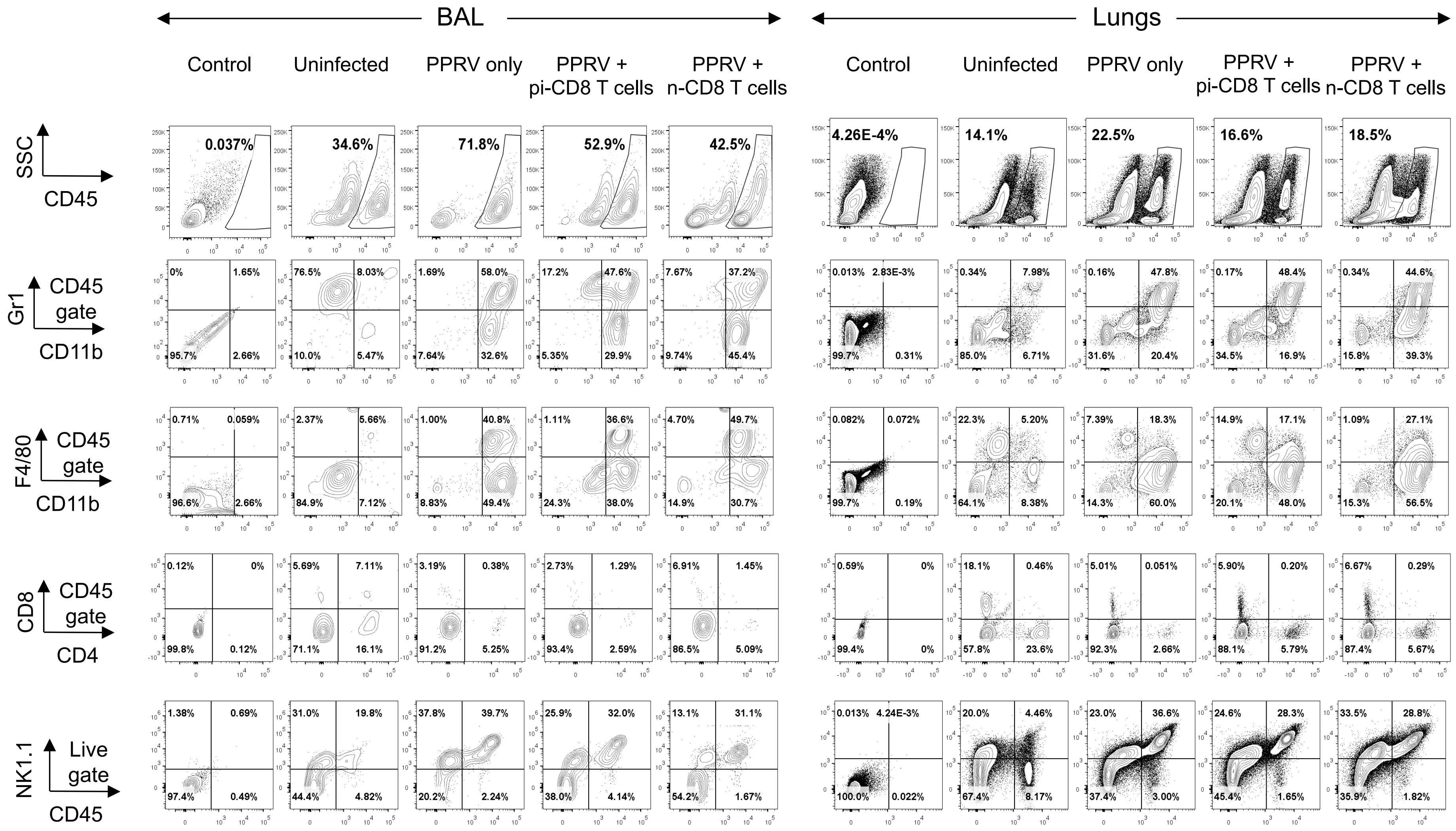


Figure 8. Representative FACS plots show the distribution of immune cells in single cell suspensions prepared from BAL and lungs of of PPRV infected IFNR KO mice transferred with WT CD8⁺ T cells collected from naïve or the previously PPRV infected mice. The control and recipients were infected with the $10^4$ of PPRV via intranasal route and scarified on 6dpi. A. The cellular analyses were performed in single cell suspension of BAL and lungs. Indicated markers are shown in the respective FACS plots

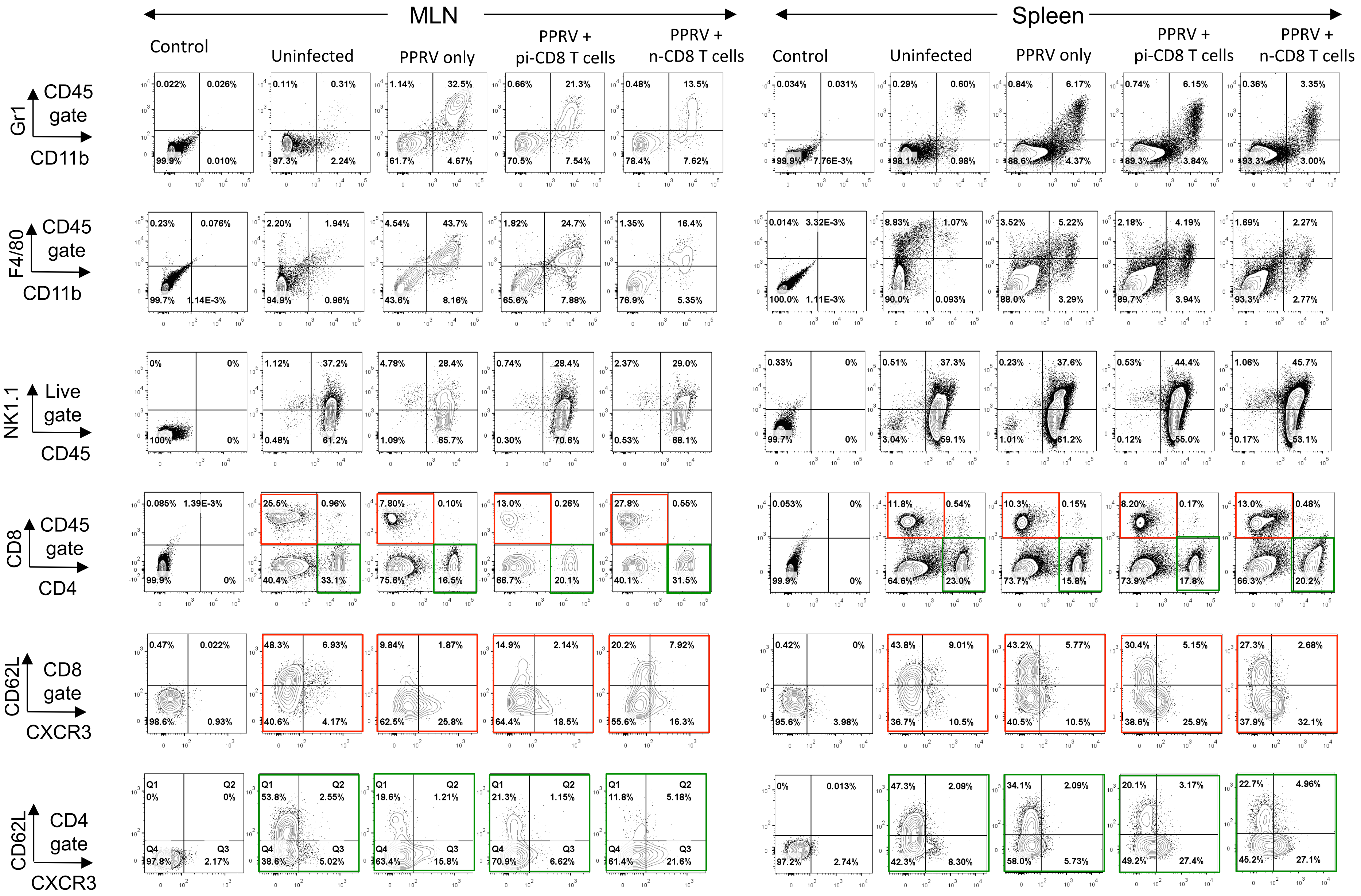


Figure S9. Representative FACS plots show the distribution of immune cells in single cell suspensions prepared from MLN and spleen of of PPRV infected IFNR KO mice transferred with WT CD8$^{+}$ T cells that were collected from naïve or the previously PPRV infected mice. The control and recipients were infected with the $10^4$ of PPRV via intranasal route and scarified on 6dpi.

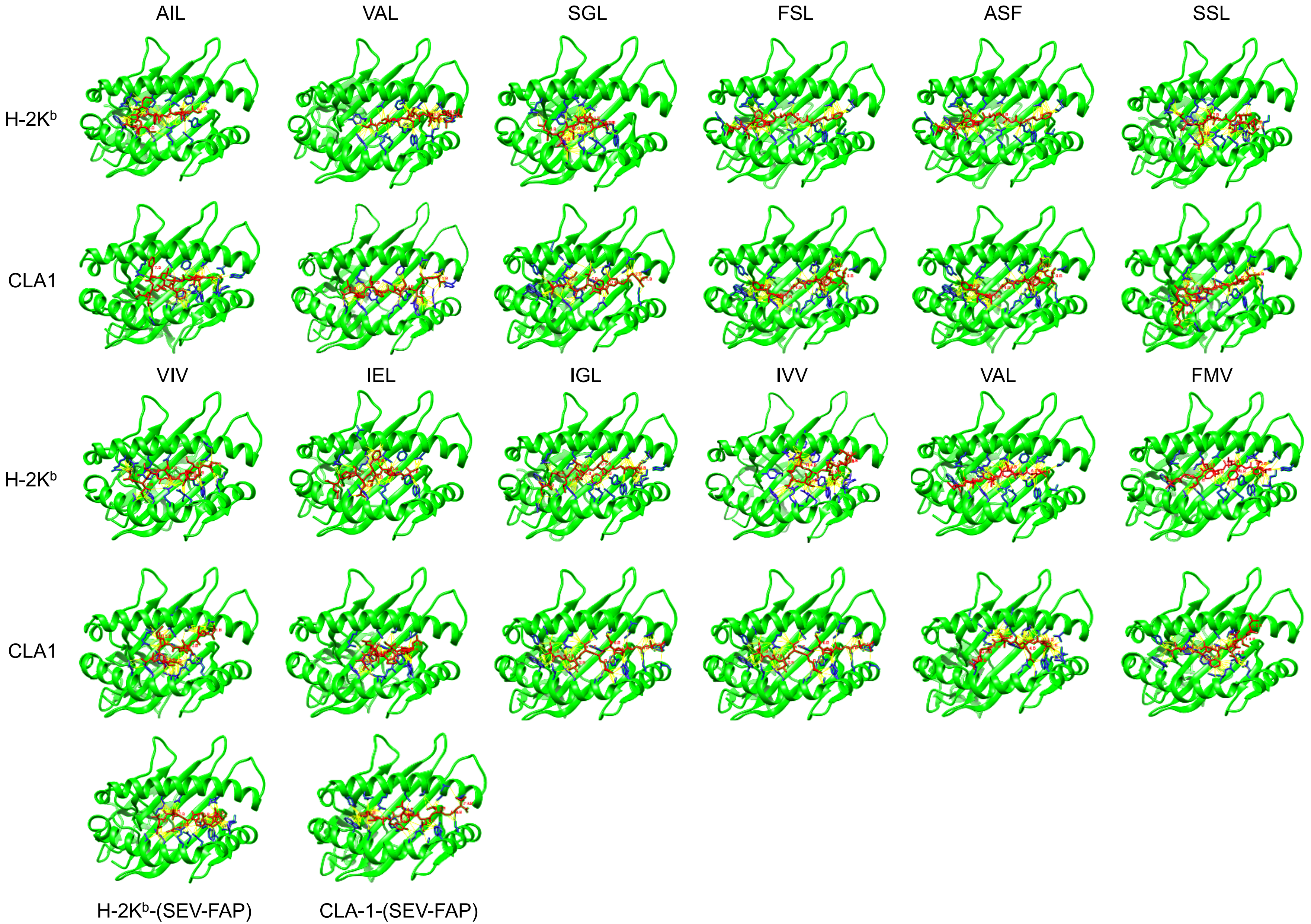


Figure S10. Molecular docking of predicted PPRV peptides for class I MHC molecule of mouse (H-2K$^b$) and goat (CLA-1). Top view of MHC-I (alpha chain) is shown green, PPRV peptide in red and interacting residues of MHC-I are shown in blue, while interactions are shown in yellow.

Table S1: A list of top scoring H-2K$^b$ restricted nonameric peptides

**Nucleocapsid**

| Peptide | Length | Score |
|---|---|---|
| ASFILTIKF | 9 | 0.18398 |
| SSIITRSRL | 9 | 0.15426 |
| FSAGAYPLL | 9 | 0.06558 |
| SIITRSRLL | 9 | 0.01413 |
| SLRRFMVSL | 9 | -0.04424 |
| STIESLMNL | 9 | -0.17957 |
| SLMNLYQQL | 9 | -0.24323 |
| INGSKLTGV | 9 | -0.32335 |
| ATLLKSLAL | 9 | -0.36699 |
| VMISMLSLF | 9 | -0.45081 |

**Matrix**

| Peptide | Length | Score |
|---|---|---|
| ANAVAFNFL | 9 | 0.26548 |
| IVVRRTAGL | 9 | 0.20524 |
| VAFNFLVTL | 9 | 0.19257 |
| FMYLFLLGV | 9 | 0.10164 |
| TFMVHVGNF | 9 | 0.07972 |
| CNAVNLVPL | 9 | 0.06586 |
| LVFYNNTPL | 9 | 0.04817 |
| VFYNNTPLS | 9 | 0.00669 |
| ECFMYLFLL | 9 | -0.06042 |
| VVYMSITRL | 9 | -0.15072 |

**Fusion**

| Peptide | Length | Score |
|---|---|---|
| SGGDFLAIL | 9 | 0.24766 |
| VAILTFLYL | 9 | 0.16852 |
| AILTFLYLF | 9 | 0.12942 |
| FLYLFPNAV | 9 | 0.1086 |
| VSLGLVTLI | 9 | 0.08484 |
| FGGNMYIAL | 9 | -0.03481 |
| SVYLHKIDL | 9 | -0.05858 |
| IAYPTLSEI | 9 | -0.06612 |
| SVHRMSCEL | 9 | -0.25115 |
| ALYPMSPLL | 9 | -0.35493 |

**Heamagglutinin**

| Peptide | Length | Score |
|---|---|---|
| IEHIFESPL | 9 | 0.20657 |
| VYYIYDTGL | 9 | 0.20256 |
| VILDRERLV | 9 | 0.20057 |
| VIERPYILL | 9 | 0.17614 |
| IGLVRDLGL | 9 | 0.11966 |
| VTRAHFSEL | 9 | 0.11675 |
| SSYYYPVRL | 9 | 0.04612 |
| VITSVFGPL | 9 | 0.01865 |
| VMFLSLIGL | 9 | -0.01258 |
| VLLVMFLSL | 9 | -0.12888 |

**Phosphoprotein**

| Peptide | Length | Score |
|---|---|---|
| SIMIAIPGF | 9 | 0.25074 |
| SEYEYEDDL | 9 | 0.22188 |
| QVQRYYVYS | 9 | 0.04008 |
| ISGATQAVL | 9 | 0.03627 |
| VYLSPEDNL | 9 | -0.07168 |
| LNIDHKDYL | 9 | -0.08942 |
| QAYHVNKGL | 9 | -0.09674 |
| SILLLKGEV | 9 | -0.14146 |
| INQSCSPAI | 9 | -0.3988 |
| RSIIKSSKL | 9 | -0.45423 |

**Large polymerase**

| Peptide | Length | Score |
|---|---|---|
| VFYLTFELV | 9 | 0.21366 |
| TQYVFYLTF | 9 | 0.16418 |
| IQYFRESLL | 9 | 0.11515 |
| VAILEYSGI | 9 | 0.00624 |
| TNFIYQQGM | 9 | -0.01868 |
| SSFDPYNMI | 9 | -0.06202 |
| LAYPRYSNF | 9 | -0.10884 |
| YNYLRCQPI | 9 | -0.11695 |
| VIDQRYSEL | 9 | -0.14356 |
| LNYLNMSRL | 9 | -0.29334 |

Table S2: Characteristics of PPRV peptides binding with Class I MHC of mice (H-2K[b])

| S.No. | Peptide | Derived from | Sequence | Docking Parameter's | | | | | |
|---|---|---|---|---|---|---|---|---|---|
| | | | | I | II | III | IV | V | VI |
| **1** | **AIL** | **Fusion Protein** | **AILTFLFLL** | **-191.11** | **Yes** | **Yes** | **No** | **Yes** | **Yes** |
| 2 | VAL | Fusion Protein | VAILTFLFL | -209.927 | Yes | No | No | No | Yes |
| 3 | SGL | Fusion Protein | SGGDFLAIL | -179.766 | No | No | Yes | No | No |
| **4** | **FSL** | **Nucleocapsid Protein** | **FSAGAYPLL** | **-220.2** | **Yes** | **Yes** | **No** | **No** | **Yes** |
| 5 | ASF | Nucleocapsid Protein | ASFILTIKF | -194.624 | Yes | No | Yes | Yes | Yes |
| 6 | SSL | Nucleocapsid Protein | SSITTRSRL | -187.869 | No | No | Yes | Yes | No |
| 7 | VIV | Matrix Protein | VILDRERLV | -160.121 | No | No | No | No | No |
| 8 | IEL | Hemagglutinin Protein | IEHIFESPL | -173.187 | No | No | Yes | No | No |
| **9** | **IGL** | **Hemagglutinin Protein** | **IGLVRDFGL** | **-176.27** | **Yes** | **Yes** | **Yes** | **Yes** | **Yes** |
| 10 | IVV | Matrix Protein | IVVRRTAGV | -194.463 | Yes | No | Yes | No | Yes |
| 11 | VATL | Matrix Protein | VAFNILVTL | -208.709 | Yes | No | Yes | No | No |
| **12** | **FMV** | **Matrix Protein** | **FMYLFLLGV** | **-243.288** | **Yes** | **No** | **Yes** | **No** | **Yes** |
| 13 | SEV | Sendai Virus | FAPGNYPAL | -233.646 | Yes | Yes | Yes | Yes | Yes |

I. Docking energy scores, II. N-terminus of the docked peptides should be deeply embedded into H2K[b] while C- terminal is held by salt-bridge near the surface of H-2K[b]. III. Residues 4 or 5 of the docked peptides may bulge out of the groove and do not make Vander Waal's contact with H-2K[b]. IV. Residue 5 or 6 are most probable primary anchor residue V. Residues at position 8 or 9 may act as secondary anchors. VI. Polar interactions (H-bonds) at peptide termini with H-2K[b] stabilize its docking.

Table S3: Characteristics of PPRV peptides binding with Class I MHC of goat CLA-1

| S.No. | Peptide | Derived from | Sequence | Docking Parameter's | | | | | |
|---|---|---|---|---|---|---|---|---|---|
| | | | | I | II | III | IV | V | VI |
| **1** | **AIL** | **Fusion Protein** | **AILTFLFLL** | **-220.883** | **Yes** | **Yes** | **Yes** | **No** | **Yes** |
| 2 | VAL | Fusion Protein | VAILTFLFL | -234.772 | No | Yes | No | Yes | Yes |
| 3 | SGL | Fusion Protein | SGGDFLAIL | -193.167 | Yes | No | No | Yes | Yes |
| **4** | **FSL** | **Nucleocapsid Protein** | **FSAGAYPLL** | **-234.857** | **Yes** | **Yes** | **Yes** | **No** | **Yes** |
| 5 | ASF | Nucleocapsid Protein | ASFILTIKF | -237.75 | Yes | Yes | No | No | Yes |
| 6 | SSL | Nucleocapsid Protein | SSITTRSRL | -206.182 | No | No | No | Yes | No |
| 7 | VIV | Matrix Protein | VILDRERLV | -175.016 | No | No | No | No | No |
| 8 | IEL | Hemagglutinin Protein | IEHIFESPL | -211.283 | No | Yes | Yes | No | No |
| **9** | **IGL** | **Hemagglutinin Protein** | **IGLVRDFGL** | **-221.898** | **Yes** | **Yes** | **Yes** | **No** | **No** |
| 10 | IVV | Matrix Protein | IVVRRTAGV | -204.106 | No | Yes | Yes | Yes | Yes |
| 11 | VATL | Matrix Protein | VAFNILVTL | -228.412 | No | No | Yes | No | Yes |
| **12** | **FMV** | **Matrix Protein** | **FMYLFLLGV** | **-262.773** | **Yes** | **Yes** | **Yes** | **No** | **No** |
| 13 | SEV | Sendai Virus | FAPGNYPAL | -258.567 | Yes | Yes | Yes | Yes | Yes |

I. Docking energy score, II. N terminus of the docked peptides should be deeply embedded into CLA-1 while the C- terminus is held by salt-bridge near the surface of CLA-1. III. Residue 4 or 5 of the docked peptides may bulge out of groove and do not make Vander-Waal's contacts with CLA-1. IV. Residue 5 or 6 are the most probable primary anchor residue V. Residues at position 8 or 9 may act as secondary anchors. VI. Polar interactions (H-bonds) at peptide termini with CLA-1 stabilize the docking.
